# Uncovering Sex Differences in the *Drosophila* Ventral Nerve Cord Through Connectome Alignment

**DOI:** 10.64898/2026.06.14.732053

**Authors:** Arie Matsliah, Christopher K Salmon, Alexander Shakeel Bates, Helen H Yang, Daniel D Lee, Lawrence K Saul, Benjamin Silverman, Jay Gager, Szi-Chieh Yu, Kyle P Willie, Austin T Burke, Ryan Willie, Doug Bland, Marissa Sorek, Celia David, Amy R Sterling, The BANC-FlyWire Consortium, Wei-Chung Allen Lee, H Sebastian Seung, Mala Murthy

## Abstract

There are now multiple *Drosophila* connectomes available, comprising both its brain and ‘spinal’ cord - comparing these connectomes requires first classifying neurons into cell types. Existing approaches for cell typing typically require extensive manual curation. Here we present a new method that automatically aligns connectomes via network topology alone. Using two complete male ventral nerve cord (VNC) connectomes as references, we assign cell types to ∼13,000 neurons intrinsic to the female VNC, and automatically identify sex-specific and sexually dimorphic cell types. We not only provide a comprehensive census of cell types across male and female nerve cords, but we uncover connectivity differences that underlie differences in function. We focus on circuits underlying song production in males and oviposition behaviors in females, and investigate the counterparts of these circuits across sexes. Our automated methods and analyses provide insights into sex differences in circuits that connect the brain and body, and pave the way for comparative connectomics at scale.

## Introduction

Recent advances in electron microscopy (EM), machine learning-based image segmentation, and technologies for proofreading neural reconstructions have collectively made possible the generation of multiple synapse-resolution wiring diagrams, or connectomes, for the *Drosophila melanogaster* model system. These include a hemibrain connectome (Scheffer et al. 2020), a whole-brain connectome (Dorkenwald et al. 2024; Schlegel et al. 2024), a ventral nerve cord connectome (Takemura et al. 2023), and, more recently, central nervous system connectomes (Bates et al. 2025; Berg et al. 2025). These comprehensive resources open new avenues for comparative connectomics, systematic comparisons of connectivity across individuals, to uncover biologically meaningful differences. Understanding how circuit architecture varies between individuals, sexes, and species can reveal which neuronal features are conserved and which are specialized. For example, discovering that the same circuit is preserved across individuals would suggest strong developmental and functional constraints, indicating that the computation the circuit performs may be general. In contrast, finding that a circuit differs (through changes in connectivity strength, neuron number, or synaptic partner identity) could reveal mechanisms of variation and plasticity (when comparing individuals), sexual dimorphism (when comparing sexes), or adaptive specialization (when comparing different species).

A key component in comparative connectomics (Costa 2025) is the ability to accurately align neurons and circuits across different datasets. Neurons within a connectome can be grouped into cell types based on developmental birth order, gene expression, morphological similarity, and connectivity (Bates et al. 2019). Such grouping is critical not only for compressing a dataset into its functional units to facilitate analysis (for example, the *Drosophila* whole-brain connectome contains ∼140,000 neurons which can be grouped into ∼9,000 cell types (Dorkenwald et al. 2024; Schlegel et al. 2024; Matsliah et al. 2024)), but also for making comparisons between datasets. The challenge is establishing homology: reliably determining which individual neuron in a new dataset corresponds to an established cell type in a reference dataset. Well-defined cell types should largely be conserved across individuals, but the fine-scale morphology and connectivity of neurons within a cell type can vary subtly, making it exceedingly difficult to find the true, morphologically and topologically equivalent partner across different connectomes without extensive manual expert curation. For example, comparing the *Drosophila* hemibrain connectome (∼20K neurons from one female fly) (Scheffer et al. 2020) with the *Drosophila* whole-brain connectome from a second female fly (Zheng et al. 2018) relied on semi-manual cell type alignment (Schlegel et al. 2024). Once aligned, it was possible to assess the extent of inter-individual variability in connectivity between the same cell types in the two different female brains.

Here, we introduce a new method for connectome alignment and apply it to matching cell types in three fly ventral nerve cord (VNC) connectomes. These datasets are the Male Adult Nerve Cord (MANC) (Takemura et al. 2024), the Male Central Nervous System (Male CNS) (Berg et al. 2025), and the female Brain and Nerve Cord (BANC) (Bates et al. 2025). In addition to matching cell types, we extend this method to automate the discovery of sexually dimorphic and sex-specific neurons in the BANC VNC and MANC connectomes. Sex differences in the *Drosophila* nervous system are well-documented and known to underlie differences in social and reproductive behaviors (e.g. (von Philipsborn et al. 2011; Asahina et al. 2014; Wang, Wang, Forknall, Parekh, et al. 2020)). Sex differences arise through the effects of the transcription factors Fruitless (*Fru*) and Doublesex (*Dsx*) (Demir and Dickson 2005; Rideout et al. 2010).

Here we define sexually dimorphic and sex-specific neurons based on connectivity; however, ultimately molecular and functional characterization of individual cell types will be required for confirmation. Recent work has similarly identified sex different neurons in female and male brain connectomes (Deutsch et al. 2025; Berg et al. 2025) through both EM-to-light microscopy (LM) and EM-to-EM comparisons. While subsets of sex-specific cell types have been identified through careful EM/LM matching and their circuitry analyzed within the MANC connectome (Lillvis et al. 2024; Stürner et al. 2025), to date there has been no comprehensive comparison between the connectomes of male and female ventral nerve cords. This presents a major gap in our understanding of the sex differences in circuitry that connect the brain and body, and it is the one we address in this study.

We find that sexually dimorphic (present in both males and females but with differences in connectivity) and sex-specific intrinsic neurons of the VNC are both strongly interconnected and concentrated in the wing tectulum and the abdominal neuromere. Our analysis of the male VNC connectome enabled expansion of the male song circuits. While females do not generate courtship songs, they nonetheless have retained a number of sexually dimorphic song neurons. Based on the remaining circuitry, most of these are likely to be vestigial, but not all. In the abdominal neuromere (ANm), we focus on the targets of sexually dimorphic descending neurons with known functions in females: DNp13 (important for ovipositor extrusion, a rejection behavior) and oviDN (important for egg laying). We find that these descending neurons diverge significantly in their targets within the ANm between males and females. While female DNp13 differentially targets motor outputs, male DNp13 targets putative neurosecretory cells that project more broadly to the periphery. On the other hand, the one dimorphic oviDN, oviDNx, targets neurosecretory cells in both sexes, though likely with sex-different outcomes.

## Results

### A New Network Alignment Method for Comparative Connectomics

Current comparative connectomics pipelines typically operate through a phased, morphology-first framework, with the assumption that morphological, spatial, or transcriptomic information provides the primary scaffold for cell identity (Schlegel et al. 2024; Stürner et al. 2025; Berg et al. 2025; Cachero et al. 2010, 2025). There are only two species (so far) for which there exist multiple connectome datasets, and for which comparisons are possible: *C. elegans* and *D. melanogaster*. In *C. elegans*, cell types are largely identified by the invariant position of the soma (Cook et al. 2025). In *Drosophila*, cell body position is not sufficiently stereotyped, and the morphology of neuronal arbors has been used for typing cells (Scheffer et al. 2020; Schlegel et al. 2024). Matching cells by morphology typically involves non-rigid spatial transformations (registering the datasets to a common coordinate space), to facilitate comparisons with existing connectomes (Schlegel et al. 2024) or LM (light microscopy) data (Deutsch et al. 2025; Stürner et al. 2025).

Morphology-first methods inherently face challenges arising from subtle variation in arbor shape caused by both reconstruction error and biological variability (Dorkenwald et al. 2022). Although reconstruction errors can affect connectivity, topology-based alignment is more robust because it decouples alignment from physical space. Specifically, morphology-based methods such as NBLAST (Costa et al. 2016) are not applicable to cell types with variable spatial positions, such as those in the optic lobes (Schwartzman et al. 2026). More generally, connectivity reflects a global topological property of the circuit rather than individual neuronal geometry, hence it is less sensitive to localized errors. This approach nevertheless requires sufficiently complete and densely reconstructed regions, though not necessarily a whole connectome.

We leveraged the Alternating Continuous and Discrete Combinatorial (ACDC) optimization technique (Lee et al., n.d.) behind the winning graph matching algorithm from a recent data challenge hosted by FlyWire (<u>codex.flywire.ai/app/vnc_matching_challenge</u>). This competition was created specifically to identify the best-performing algorithms for our connectome matching objective function - *weighted intersection* (formalized and justified in the Methods section). The challenge featured dozens of participants, including experts in connectome graph matching methods like FAQ (Vogelstein et al. 2015) and SANA (Mamano and Hayes 2017) (see details in Methods).

ACDC alternates between two complementary ways of searching for a good match between weighted, directed graphs such as connectomes. The AC step performs continuous optimization, where the discrete matching problem is relaxed to allow gradient-based Frank-Wolfe optimization method (Frank and Wolfe 1956), while the DC step performs discrete combinatorial search (e.g. swapping pairs of nodes) to increase the objective score. By repeatedly alternating between these two steps, the continuous updates help escape local optima reached by the discrete search, while the discrete updates refine the solution to produce a valid permutation that maximizes the final matching score.

One caveat is that ACDC requires equally sized connectomes as input and produces a strict one-to-one alignment between neurons. While suitable for a graph matching contest, this assumption is biologically unrealistic because homologous cell types frequently differ in abundance across hemispheres and individuals (Schlegel et al. 2024). For example, 1,174 of the 3,425 cell types shared between MANC and Male CNS (34%) contain different numbers of neurons in the two datasets (Fig. S2f). To address this limitation, we use the ACDC alignment and its coordinate-wise confidence scores to define a continuous connectivity similarity metric between any pair of neurons across datasets (Fig. 1a; Methods). This allows neurons to be reassigned when their connectivity is more similar to a neuron other than their initial ACDC match, thereby relaxing the rigid one-to-one correspondence imposed by graph matching.

**Fig. 1.**
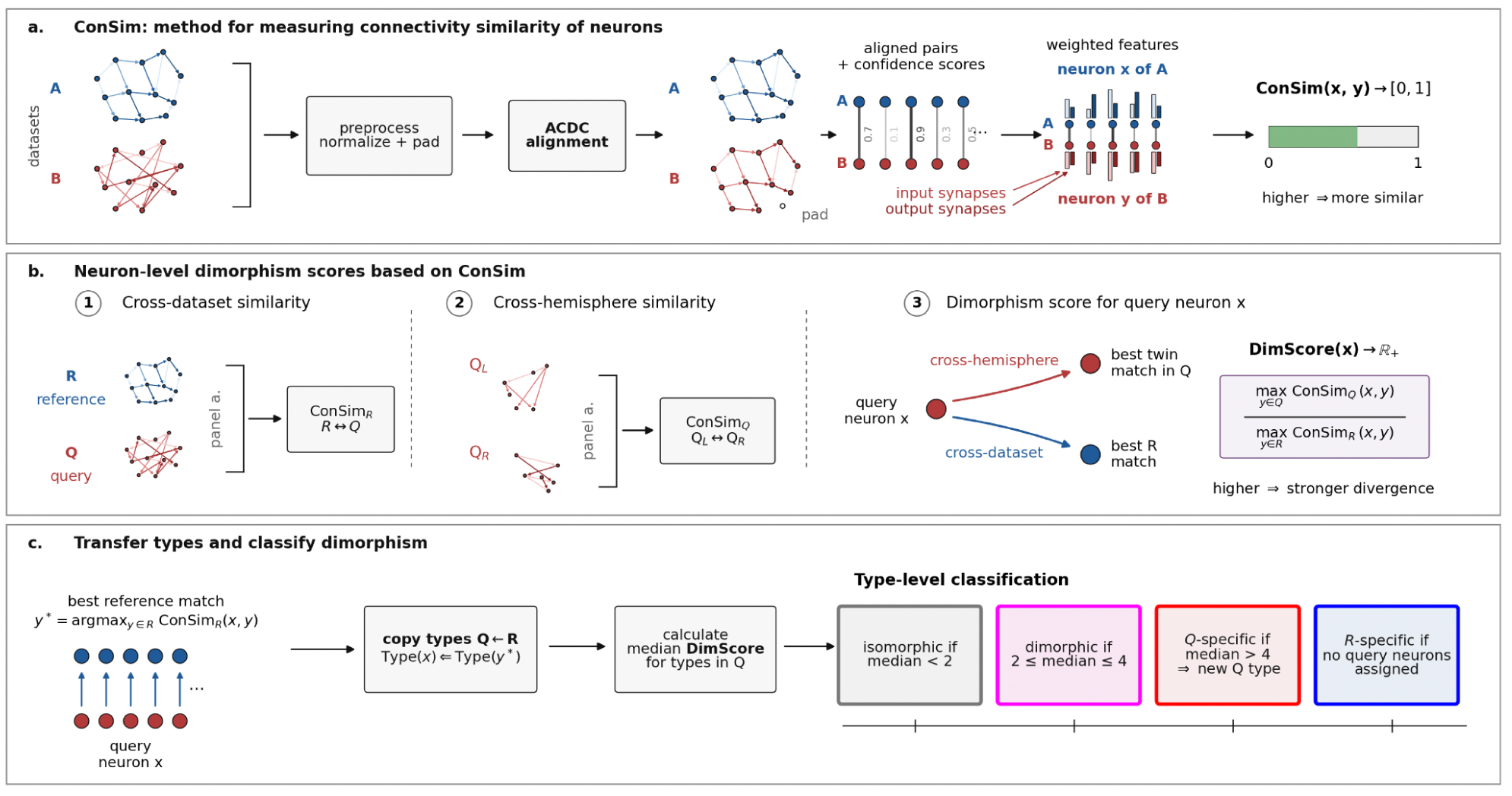
Connectivity-based alignment, cell type transfer and dimorphism identification. (a) Construction of the connectivity similarity metric (ConSim). Two connectomes are first preprocessed by normalizing synaptic weights and adding padding nodes to equalize graph size. The connectomes are then aligned using the ACDC graph matching algorithm, producing a one-to-one correspondence between neurons together with alignment confidence scores. These confidence scores are used to construct dynamically weighted connectivity feature vectors, in which each coordinate represents the number of input or output synapses associated with an aligned partner neuron. Connectivity similarity between neurons x and y is computed by comparing their weighted feature vectors, yielding a similarity score ConSim(x,y) ranging from 0 to 1. See additional details in Fig. S1. (b) Computation of neuron-level dimorphism scores. ConSim is applied independently to two comparisons. First, neurons in a query connectome (Q) are compared against neurons in a reference connectome (R), producing cross-dataset similarity scores. Second, neurons in the left and right hemispheres of the query connectome (Q_L_ and Q_R_) are compared, producing cross-hemisphere similarity scores. For each query neuron x, the neuron dimorphism score (DimScore) is defined as the ratio between the highest similarity to a neuron in the opposite query hemisphere and the highest similarity to a neuron in the reference connectome. High DimScore values indicate that a neuron is substantially more similar to its within-query counterpart than to any neuron in the reference dataset, consistent with stronger divergence between the two connectomes. (c) Cell-type transfer and dimorphism classification. Cell-type annotations are transferred from the reference connectome to the query connectome by assigning each query neuron the type of its most similar reference neuron. Dimorphism scores are then aggregated at the cell-type level by computing the median DimScore across all neurons assigned to a given type. Types are classified as isomorphic (median DimScore < 2), dimorphic (2 ≤ median DimScore ≤ 4), query-specific (median DimScore > 4), or reference-specific when no query neurons are assigned to the type.

The same framework can be applied to comparisons across individuals, hemispheres, and sexes (Fig. 1b-c). In its current implementation, ACDC can practically be applied to datasets containing approximately 20,000–25,000 neurons. This limitation arises from the use of explicit connectivity matrices during global permutation optimization, which becomes prohibitively memory intensive for larger graphs. The neuron populations analyzed in this study - INs, neck ANs/DNs (ascending/descending neurons), and MNs (VNC motor neurons) - fit within this limit and can be aligned jointly. In contrast, alignment of an entire adult fly connectome would currently require a piecewise strategy, for example by aligning individual regions separately and enforcing consistency between successive alignment steps through additional constraints that preserve the identities of neurons shared between regions.

### Application of Network Alignment to Cell Typing and Identification of Sex Differences in Connectivity

We applied the new framework to identify cell types and categorize sexually dimorphic and sex-specific neurons in the BANC VNC. Unlike ANs, DNs, and MNs, for which many sexually dimorphic and sex-specific types have already been identified and manually characterized, sex differences among VNC intrinsic neurons (INs) remain largely unexplored. ANs, DNs, and MNs were included in the alignment and used to calibrate the classification framework against known dimorphisms, but subsequent discovery analyses focused on INs.

We first aligned the BANC VNC (query) to the MANC VNC (reference), including neck neurons (ANs and DNs), motor neurons (MNs), and intrinsic neurons (INs), while excluding sensory neurons (SNs) because their abundance differs substantially between datasets (Fig. 2a,b). For each neuron in the query dataset (BANC), we compared similarity to its cross-hemisphere twin and similarity to its closest cross-dataset match (Fig. 2c). We also performed the reciprocal analysis using BANC as the reference and MANC as the query, allowing DimScore distributions to be evaluated from the perspective of MANC neurons (Fig. 2d).

**Fig. 2.**
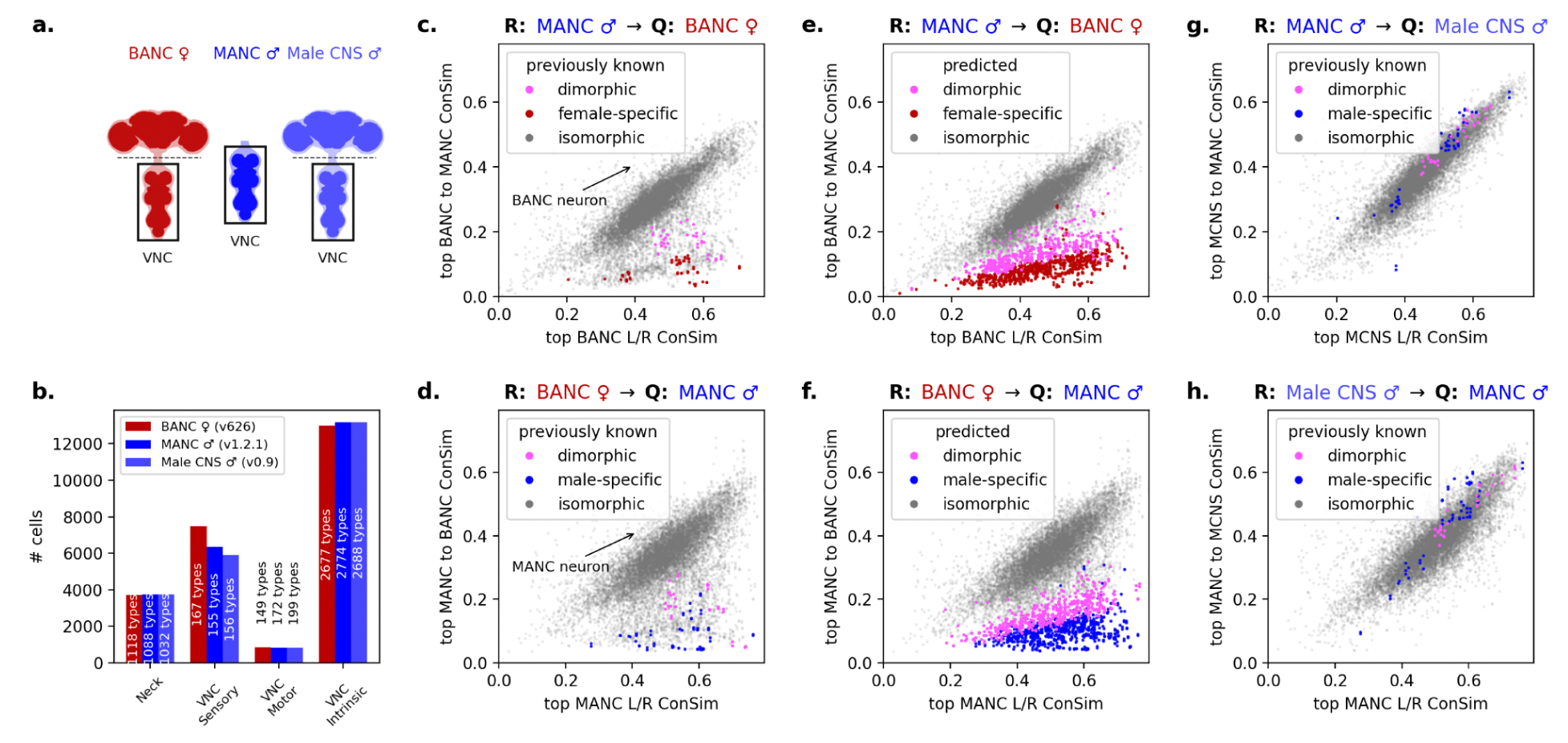
Connectivity-based dimorphism classification. **(a)** The three datasets being compared. **(b)** Composition of the VNC datasets. Bars indicate the number of neurons within four major classes: neck neurons (ANs and DNs), VNC sensory neurons (SNs), VNC motor neurons (MNs), and VNC intrinsic neurons (INs). Numbers inside the bars indicate the corresponding number of annotated cell types. **(c,d)** Validation using previously characterized (‘known’) dimorphic and sex-specific neurons. Scatter plots show neuron-level connectivity similarity scores for comparisons between MANC and BANC. The x-axis indicates the highest cross-hemisphere connectivity similarity (top L/R ConSim), while the y-axis indicates the highest cross-dataset connectivity similarity (top ConSim to the reference connectome - reference connectome is MANC in c and BANC in d). Previously characterized dimorphic, male-specific, and female-specific neuron types identified through literature review (Table 1) are highlighted and used to calibrate classification thresholds. **(e,f)** Application of DimScore-based classification to all neuron types in MANC (e) and BANC (f). Neurons assigned to cell types classified as sexually dimorphic or sex-specific according to the thresholds are highlighted. Remaining neurons are classified as isomorphic. These classifications are subsequently used for cell-type annotation and identification of sex-specific populations. **(g,h)** Baseline inter-individual variability estimated from comparisons between two independent male VNC connectomes, MANC and Male CNS. Previously characterized dimorphic and male-specific neuron types are highlighted. Unlike cross-sex plots, most neurons remain concentrated along the diagonal, indicating that connectivity patterns are largely preserved across individuals of the same sex and establishing a baseline against which male/female differences can be interpreted.

Although ACDC produces a one-to-one alignment between neurons, subsequent cell-type transfer was performed using ConSim (Fig. 1a). Initially each query neuron (in BANC) was assigned to a type of its highest-similarity reference neuron (in MANC). Cell types in the query dataset were then classified using the median DimScore across all neurons assigned to a type (Fig. 1c). Types with median DimScore < 2 were classified as isomorphic, types with median DimScore between 2 and 4 as sexually dimorphic, and types with median DimScore > 4 as sex-specific. Thresholds of 2 and 4 were selected as the nearest simple integer cutoffs that correctly separated the previously characterized sexually dimorphic and sex-specific neuron types used as biological ground truth (Table 1). Applying these criteria identified 205 male-specific IN types (659 cells, Fig. 2e (blue)), 190 female-specific IN types (417 cells, Fig. 2f (red)), and 144 sexually dimorphic IN types represented in both datasets (587 cells in MANC (Fig. 2e (magenta)) and 582 cells in BANC (Fig. 2f (magenta)). The identified dimorphic and sex-specific neurons span all major neuromeres of the VNC (Fig. 3f).

**Fig. 3.**
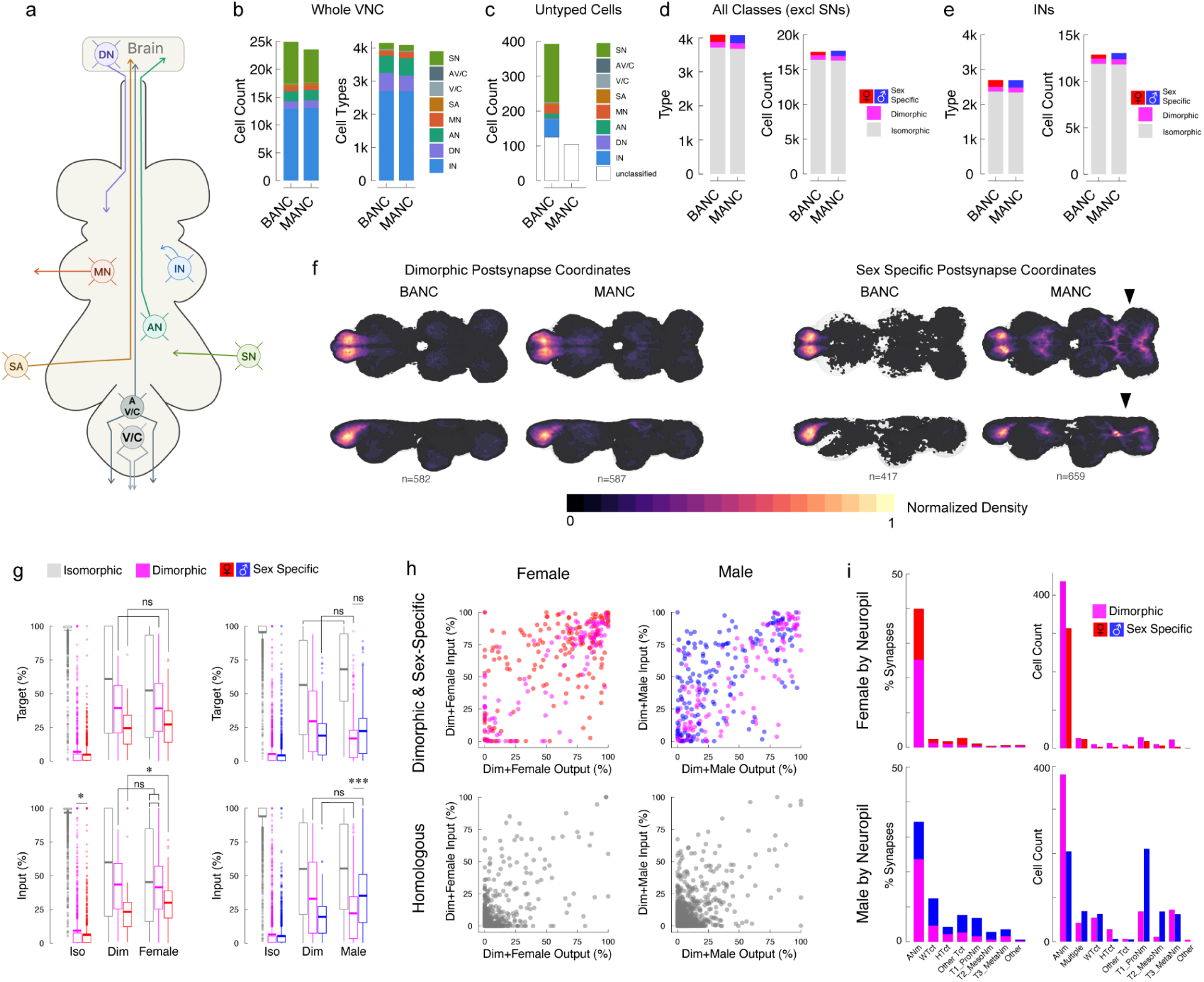
Sex-specific and dimorphic neurons in the VNC. **a)** Schematic of cell super classes in the VNC (naming conventions correspond to (Bates et al. 2025)). **b)** Comparison of the numbers of cells and cell types in each super class. **c)** Cells that have been fully or partially reconstructed in MANC and BANC VNC that are as of yet untyped. Cells without a final super class assigned are indicated by the empty portion of the bars. **d)** Counts of cells and cell types across all super classes broken down by their dimorphism. **e)** Counts of INs and IN types by dimorphism. **f)** Kernel density estimate of synapse density based on postsynapse coordinates presented in JRC2018VNCU reference brain space. Arrowheads indicate the mesothoracic triangle. **g)** Summary stats showing the proportion of isomorphic, dimorphic and sex-specific targets and inputs of other isomorphic, dimorphic, and sex-specific cell types. Each box represents the isomorphic, dimorphic, or sex-specific target or input of the category noted on the x-axis. Each data point in each bar is a cell type. Boxplot central bars indicate mean. **h)** Plots of the proportion of input and output connections that are dimorphic and sex-specific for each IN cell type. Each point represents an IN cell type. Points are colored by the cell type’s own classification: isomorphic (grey), dimorphic (magenta), or sex-specific (red/blue). **i)** Proportion of each neuropil taken up by dimorphic (magenta) and sex-specific (female red, male blue) synapses (left) and cells (right).

For cell types classified as isomorphic or sexually dimorphic, annotations were transferred directly from the corresponding MANC cell type. Consistent with the inferred classifications, we observed strong agreement in both cell-type abundance and type-to-type connectivity among isomorphic and sexually dimorphic populations (Fig. S2d,j). Types classified as sex-specific were instead grouped by cross-hemisphere connectivity similarity and assigned provisional placeholder identities with decreasing index and *_*f suffix (e.g. INXXX999_f), akin to the naming scheme in (Stürner et al. 2025).

During the course of this work, a complete male central nervous system dataset (Male CNS) became available (Berg et al. 2025), enabling an independent assessment of same-sex inter-individual variability. MANC and Male CNS originate from males of the same genotype and were reconstructed by the same group using the same techniques, therefore providing a high quality baseline of inter-individual variability in the *Drosophila* VNC. As an additional validation, we aligned the same population of neurons between MANC and Male CNS VNC (Fig. 2g with MANC as reference, Fig. 2h with Male CNS as reference), and found that in contrast to the male/female comparisons, most neurons occupied a narrow diagonal band in both directions (Fig. 2g,h), indicating that connectivity patterns are largely preserved between independent male VNCs. These observations suggest that the inferred correspondences are biologically meaningful and motivate a more systematic evaluation of the alignment framework.

### Validation of Connectome Alignment and Cell Type Assignment

We validated the connectivity-based cell typing and dimorphism classification approach in two ways: quantitative consistency and expert review against morphology-based matching.

### Consistency checks

We found that the inferred cell type composition closely matched the reference connectome, including hemisphere-specific counts (Fig. S2a), the number of instances per type (Fig. S2d), and synapse distributions between pairs of types across hemispheres (Fig. S2g). Among isomorphic and sexually dimorphic populations, we observed strong agreement in both cell-type abundance and type-to-type connectivity (Fig. S2d,j). Similar results were obtained when comparing against Male CNS, indicating that the inferred correspondences are stable across independent male connectomes (Fig. S2k,l).

### Expert review

In cases where connectivity-based and morphology-based matches disagreed, we performed manual review. First, an expert reviewer (CKS) examined 300 cases in which the top NBLAST match differed from the connectivity-based match. Both morphology and up- / downstream partners were examined in the review. The connectivity-based match was judged to be better in 62% of cases, whereas the top NBLAST match was better in 7% of cases. In 26% of cases, connectivity-based matching and NBLAST identified equally plausible matches belonging to different cell types, and in the remaining 6% of cases both approaches produced poor matches. Thus, connectivity-based matching performed as well as or better than the top NBLAST match in 88% of cases. In practice, however, cell typing rarely relies on the top NBLAST hit alone; expert annotators typically examine the top three to five NBLAST candidates before selecting a final match. When an expert reviewer (ASB) visually assessed all 2,899 INs assigned to different types by connectivity-based and NBLAST-based matching, connectivity-based matching was preferred in approximately 75% of cases (2,174/2,899), compared with approximately 22% for NBLAST and approximately 3% in which neither approach produced a satisfactory match. Together, these analyses indicate that, when morphology- and connectivity-based assignments disagree, connectivity-based matching performs as well as or better than the top morphology-based hit in 88% of reviewed cases and is preferred over expert-reviewed top-five morphology-based candidates in approximately 75% of cases.

### Comparing intrinsic neurons of female and male VNCs

Having assigned and classified IN types in the BANC VNC by dimorphism status, we next examined their anatomical distribution and connectivity. Individual cell and cell type counts are similar to the reference (MANC), with the exception of sensory neurons (SNs) (Fig. 3a-b), which are more numerous in BANC (Bates et al. 2025). A small number of cells remain untyped in both datasets (Fig. 3c). While some sexually dimorphic and sex-specific ANs and DNs have been studied (Feng et al. 2014; Wang, Wang, Forknall, Parekh, et al. 2020; Wang, Wang, Forknall, Patrick, et al. 2020), outside of the male song circuitry, e.g. (Lillvis et al. 2024), there were no sexually dimorphic INs characterized in the VNC before this work.

Together with previously characterized VNC dimorphisms (Stürner et al. 2025; Lillvis et al. 2024; Yu et al. 2010), our results substantially increase the known repertoire of sex-specific and sexually dimorphic VNC cell types, adding 190 female-specific IN types (417 cells), 205 male-specific IN types (659 cells), and 144 dimorphic IN types represented in both BANC (582 cells) and MANC (587 cells) (Fig. 3d,e). The numbers of isomorphic types are uneven in BANC and MANC due to unrectified naming conventions (2362 types BANC; 2344 types MANC). We expect that extending our network alignment approach to sensory and effector neurons will rectify this discrepancy while adding more dimorphic types to the growing inventory (for example, there are known sex-specific SNs in the legs (Kallman et al. 2015)) that are not characterized here). Many of the sexually-dimorphic and sex-specific IN types we detect are ‘new’, or never previously characterized. These include new types that likely function in the production of male courtship song (see Fig. 4 and 5) and female oviposition behaviors (see Fig. 6 and 7).

**Fig. 4.**
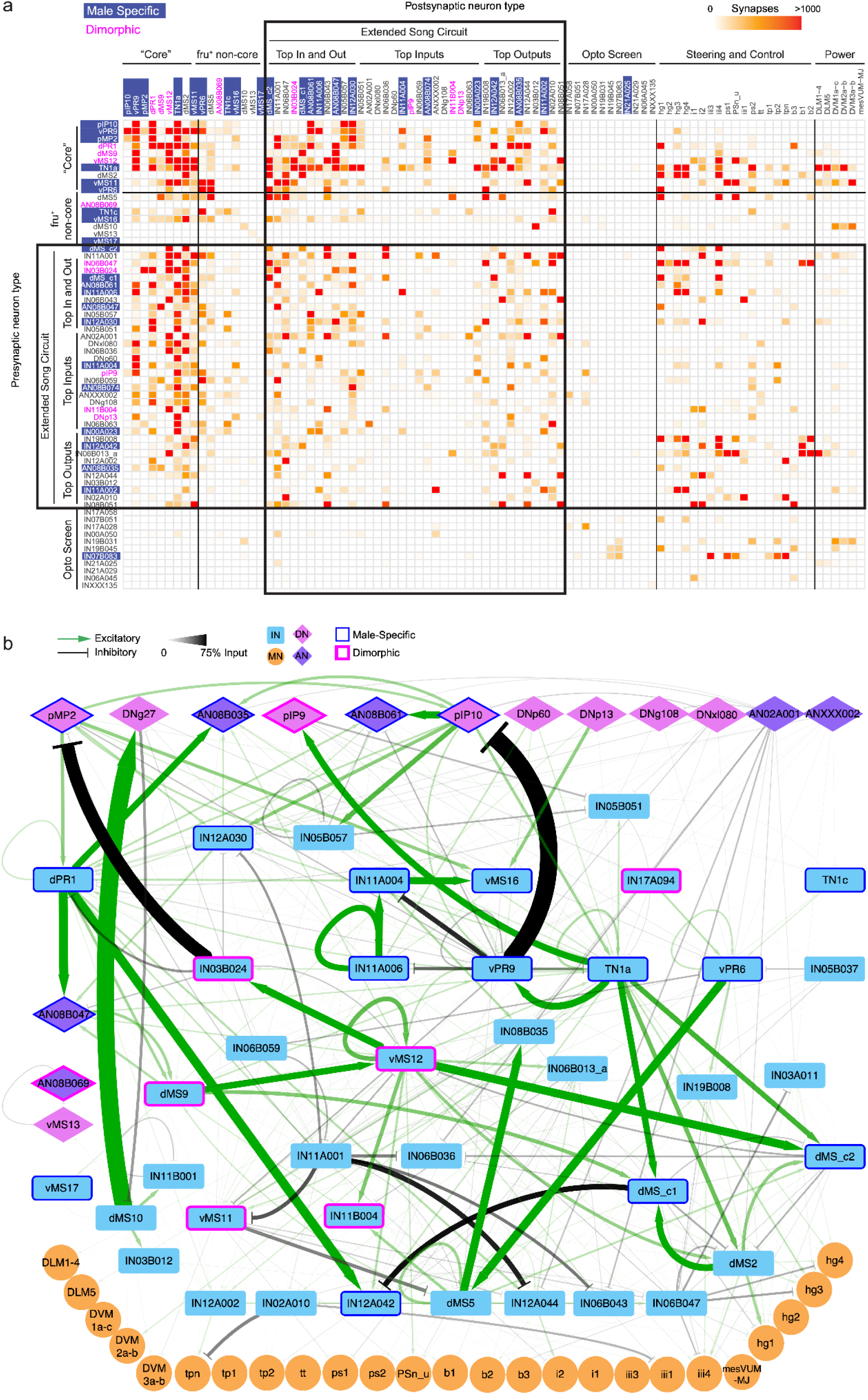
Song-Related Circuitry of the Male Wing Tectulum. **a)** Connectivity matrix showing connections between “core” song neurons; other *fru^+^* song neurons; neurons optogenetically screened for roles in song (Lillvis et al. 2024); and the extended song circuit divided into top inputs to the core, top targets of the core, and top summed input+output connections from/to the core. b) The entire proposed “extended song circuit,” including the core and the non-core *fru^+^* song neurons as defined by <u>(Lillvis et al. 2024)</u>. Edge weight corresponds to input percentage (the proportion of the target cell type’s total input made by the upstream cell type).

**Fig. 5.**
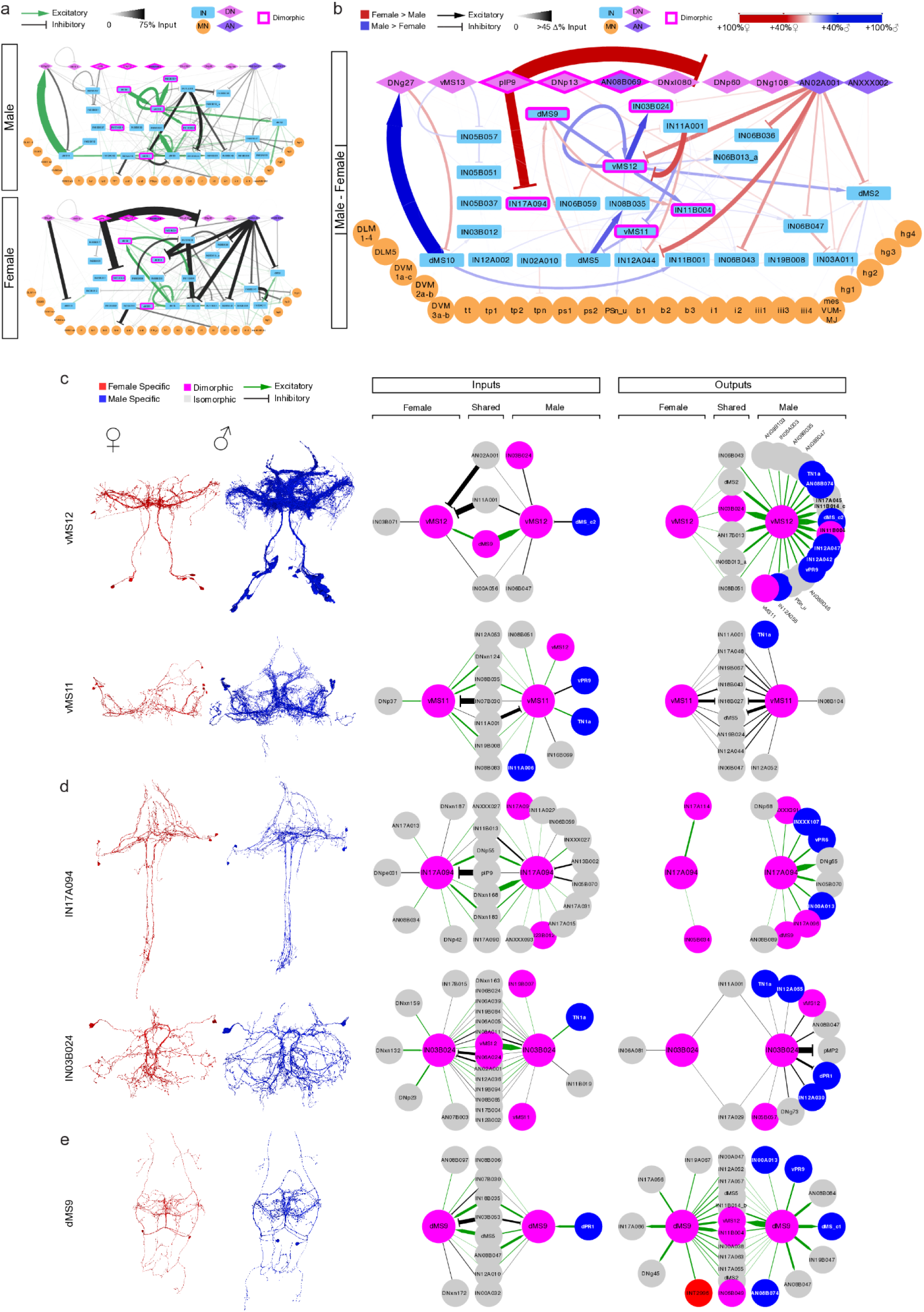
Song circuit elements that are preserved in the female. **a)** Network diagrams showing cell types of the extended song circuit that are present in both male (top) and female (bottom). b) Female-male comparative graph with edge weights set to the absolute difference between the male and female percent input made by the upstream cell type. c-e) Renders (left) and single cell type comparative graphs (right) of the dimorphic cells in the shared extended song circuit. Connections shown are thresholded at 1% input to target, with the exception of some connections to shared cell types: as long as a shared type gets >1% input from one sex, it is considered shared if it has any connection, including <1%, from or to the shared type. This is evident in the vMS12 output graph.

**Fig. 6.**
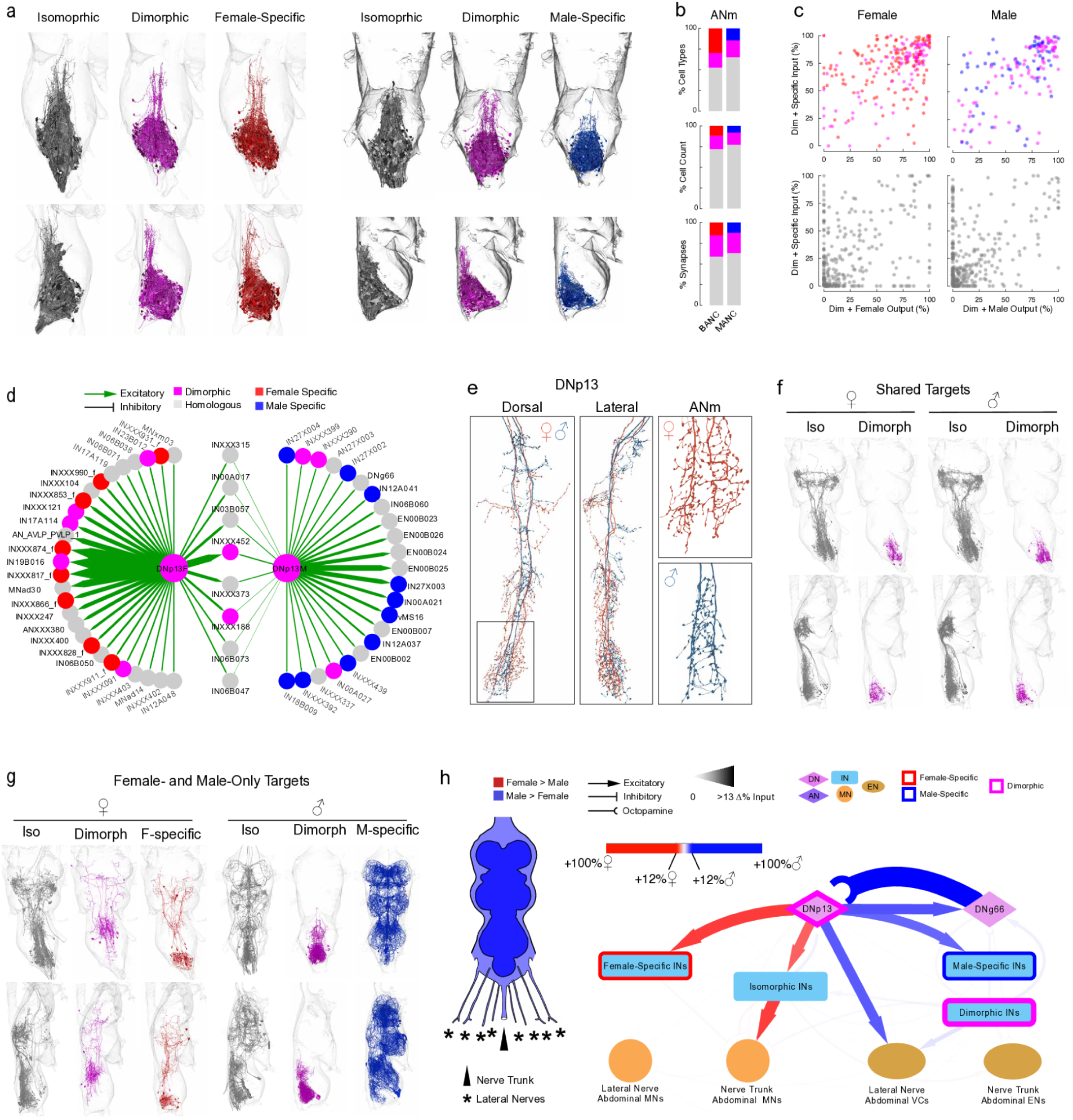
ANm cell types and DNp13 downstream circuitry in males and females. **a)** Full complement of isomorphic, dimorphic (magenta) and sex-specific (female, red; male, blue) INs in the ANm. b) Proportions of ANm cell types, cell counts, and synapses that are dimorphic or sex-specific. c) Percent input from dimorphic and sex-specific cell-types plotted against percent output to dimorphic and sex-specific neurons. Only INs of the ANm are considered (top: magenta dimorphic ANm cell types, red female-specific ANm cell types, blue male-specific ANm cell types; bottom, gray isomorphic cell types). d) Comparing DNp13 outputs in male and female VNC. Only connections accounting for >5% of target cell-type’s input synapses are shown. e) Portion of DNp13 in the VNC in BANC (red) and MANC (blue). Box at bottom left indicates the portion of DNp13 in the ANm and is shown magnified at right separately for female (top) and male (bottom). f) 3D renders of DNp13 targets shared in males and females. Male and female neurons are both presented in BANC space for ease of comparison. g) DNp13 targets present in only male or female VNC. Note that this includes isomorphic neurons that receive inputs from DNp13 in only one sex or the other. (e,f: top row, dorsal view; bottom row, lateral view with dorsal aspect facing left). h) Comparative graph with edge width equal to the absolute difference between male and female input percentage by target. MNs and VCs (ENs) are grouped by their exit nerve. INs are grouped by dimorphism. (h: the 5% threshold was applied to INs, DNs and ANs; MNs and ENs tend to have many proportionally weak inputs, and thus all connections to effectors were considered regardless of weight.)

**Fig. 7.**
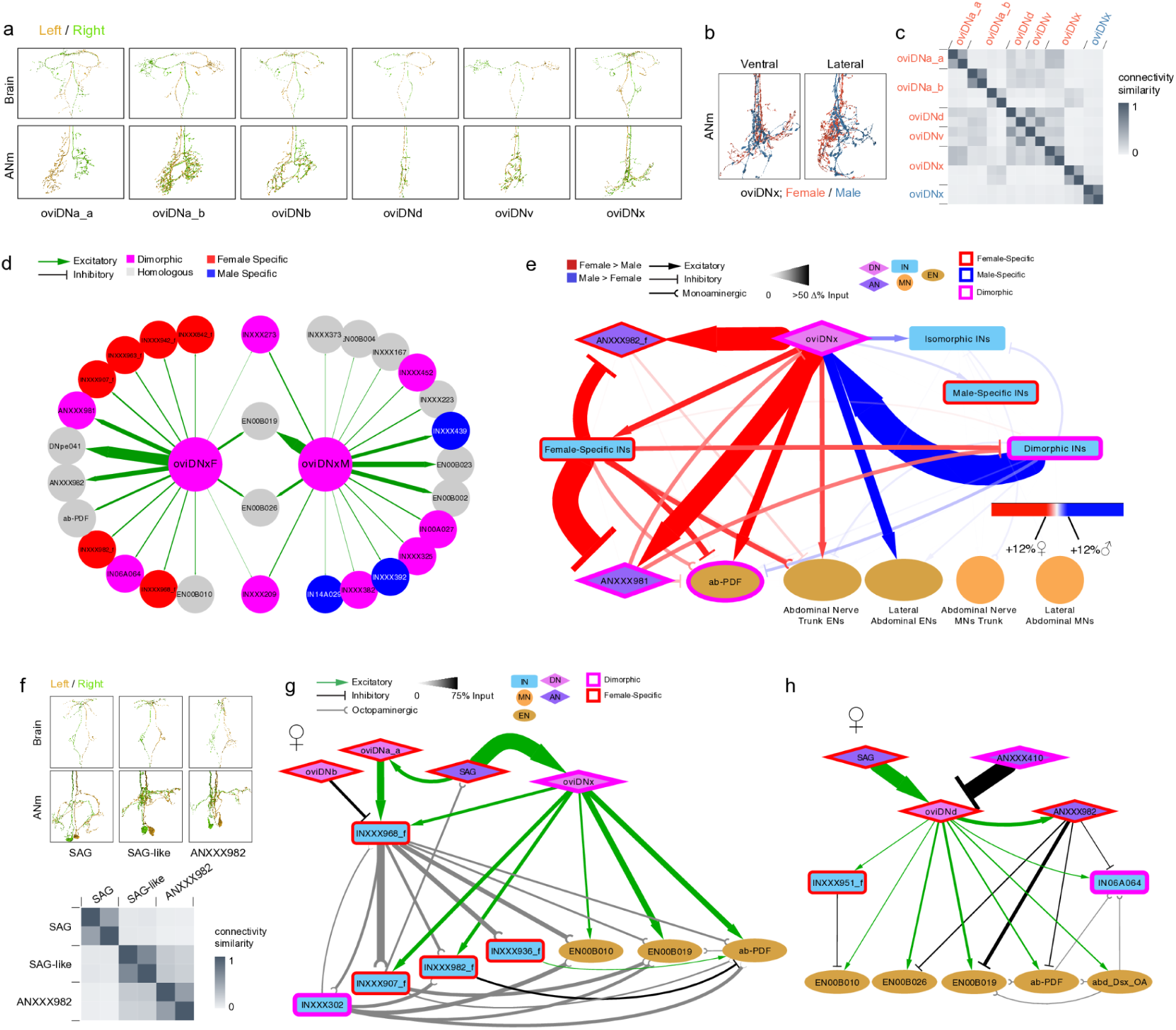
Circuitry downstream of oviDNs in the male and female ANm. **a)** Renders of all female oviDN types. b) Male and female oviDNx arbors in the ANm. c) Similarity matrix comparing all oviDN types in female and male using mean cell type ConSim values. d) Outputs of oviDNx in female (left) and male (right) ANm. Targets for which oviDNx accounts for >2% of input synapses are included. e) Comparative graph with edge width equal to the absolute difference between male and female input percentage by target. MNs and ENs other than ab-PDF are grouped by their exit nerve. INs are grouped by dimorphism category. ANs and DNs are left ungrouped. Note that male oviDNx makes no major connections with ANs or DNs in the VNC. f) Renders of key oviDNx AN targets in females. g) Broad octopaminergic modulation of ANm targets through INXXX968_f downstream of oviDNa_a. h) ANm targets downstream of oviDNd.

Prior studies indicate that sexually dimorphic circuitry is concentrated in the Abdominal Neuromere (ANm), which controls reproductive function, as well as the male wing tectulum, which controls male courtship song (Cachero et al. 2010; Yu et al. 2010). We plotted the density of synapses made by our identified dimorphic and sex-specific INs in BANC and MANC (Fig. 3f), confirming that these cells largely make connections in the expected neuropils. In particular, the “mesothoracic triangle” is specifically innervated by male-specific neurons.

It has been reported that dimorphic neurons connect more strongly with other dimorphic neurons (Stürner et al. 2025; Berg et al. 2025; Deutsch et al. 2025). We find this is also the case with the population of dimorphic VNC INs we uncovered. Both known (Fig. S3) and newly predicted sexually dimorphic INs (Fig. 3g-i) receive and make a higher proportion of connections with other dimorphic neurons. Interestingly, in the female, dimorphic and female-specific cell types both bias toward connections with dimorphic neurons (Fig. 3g left), while in the male, dimorphic neurons make more connections with dimorphic neurons and male-specific neurons make more connections with other male-specific neurons (Fig. 3g right). This may reflect differences in the wing tectulum (known to be enriched for male-specific neurons). Interestingly, some *isomorphic* types also have varying levels of dimorphic and sex-specific inputs and targets (Fig. 3g), and relative input-output plots indicate that a small minority of isomorphic types approach the levels of dimorphic and sex-specific/dimorphic connectivity present in dimorphic and sex-specific types (Fig. 3h, S3b). This observation is not unexpected because DimScore (Fig. 1b) provides a continuous measure of connectivity divergence, whereas the classification into isomorphic, dimorphic, and sex-specific types imposes discrete boundaries on an underlying continuum. Consequently, some types classified as isomorphic may nevertheless participate extensively in sex-biased circuitry while retaining sufficiently similar overall connectivity profiles to fall below the dimorphism threshold. The opposite is also true - some dimorphic and sex-specific types have few connections to other dimorphic or sex-specific types. These varying levels of dimorphic connectivity should be considered when interpreting the potential roles of these cells in sex-different behaviors.

We next computed the proportion of synapses in each neuropil associated with dimorphic and sex-specific IN types; while the ANm has a high proportion in females, there is more variability in males (Fig. 3i, S3b). Interestingly, while the wing tectulum (WTct) is known to harbour male-specific and dimorphic neurons, it is not the primary site of dimorphic and male-specific synapses outside the male ANm; many dimorphic IN types also innervate the leg neuropils (ProNm, MesoNm, MetaNm) and other neuropils as well (Fig. 3i, S3b-d).

### Sex Dimorphisms in the Wing Tectulum

During courtship, *Drosophila* males generate complex song sequences that are patterned relative to female cues (Coen et al. 2014; Roemschied et al. 2023); these songs are generated by unilateral wing vibration (Shorey 1962). Recent work has functionally characterized an array of cell types in the male VNC important for courtship song (Lillvis et al. 2024); this revealed a complex circuit responsible for the production of the two main syllables of courtship song, pulse and sine (Clemens et al. 2018; Shiozaki et al. 2024). Many of the neurons of the song circuit are known to be sexually dimorphic or sex-specific, based on comparisons with LM images and expression of Fruitless and Doublesex (Yu et al. 2010; von Philipsborn et al. 2011; Rideout et al. 2010; Shirangi et al. 2016), but no prior study has systematically compared the neurons of the wing tectulum between male and female EM datasets. We not only identify new sexually dimorphic and male-specific cell types of the song circuit, but we examine its counterpart in the female VNC where the purpose of preserved dimorphic cell types is unclear since only males sing.

We first expanded the song circuit. The original analysis focused on cell types identified through genetic screening (Lillvis et al. 2024); instead, we focused on connectivity. In MANC, we identified neurons receiving >1% of their input from a set of “core” song neurons (Lillvis et al. 2024), and/or constituting >1% of core neurons’ input (Fig. 4a,b). This “expanded” song circuit contains isomorphic, dimorphic and male-specific neurons that strongly target and are strongly targeted by the core song circuit (Fig. 4a, “Core”). With this approach we identified 35 new cell types putatively involved in courtship song (12 male-specific, 4 dimorphic, 19 isomorphic). A subset of the expanded song circuit directly targets the wing steering MNs at levels similar to the core song types (Fig. 4a, “Steering and Control”). In comparison, putative song neurons identified through optogenetic screening in (Lillvis et al. 2024) - which had only weak effects on song - make only weak connections to the wing steering MNs (Fig. 4a, “Opto Screen”). To visualize how the expanded song circuit fits into the currently known song circuit, we plotted the network graph with edge weights equal to the percent of the target cell type’s *input* accounted for by the upstream cell type (Fig. 4b). We reason that percent input best approximates the influence that an upstream cell exerts over the target cell type (in contrast with prior network diagrams of the song circuit (Lillvis et al. 2024; Stürner et al. 2025)).

One newly predicted song cell type, IN03B024, provides strong feedback inhibition to pMP2 and dPR1 and receives excitatory input from vMS12, all three of which are involved in pulse song production (Fig. 4b) (Lillvis et al. 2024). This feedback loop may be important for generating song patterns, similar to the role of feedback inhibition in walking circuits (Pugliese et al. 2026). IN03B024 allocates more of its output (79%) to dimorphic and male-specific types than any other cell type intrinsic to the wing tectulum, and receives ∼50% of its input from dimorphic and male-specific neurons (Fig. S3). That this cell type was missed previously highlights the power of using the connectome to drive hypotheses about circuit function.

We next compared the part of the song circuit present in both males and females (Fig. 5a) and generated a comparative circuit diagram showing relative changes in connection weights between male and female (Fig. 5b). Compared to circuitry known to control unisex wing behaviour, such as flight steering and wing power muscles, the portion of the male song circuitry present in both sexes has much higher male-female divergence (Fig. S5). This divergence is strong in connections involving both dimorphic and isomorphic cell types. One major difference is the influence of inhibitory neurons in the female: pIP9 (dimorphic, predicted to be glutamatergic (Bates et al. 2025)) and AN02A001 (isomorphic, also glutamatergic) exert strong influence over both dimorphic and isomorphic “song” neurons in females. Closer examination of the targets of AN02A001 shows that this cell type makes larger proportional input to dimorphic downstream neurons like vMS12 and dMS2 in the female because those cell types are missing inputs from male-specific neurons in the female (Fig. S6).

To probe the possible function of individual dimorphic song-like neurons in females, we generated single cell-type male-female comparative input and output graphs (Fig. 5c-e). A shared connection (middle columns) is defined as any cell type that receives input from both sexes, while unshared connections (left and right semicircular groupings) are present in only one sex (note that by this definition, unshared connections need not be sex-specific and vice-versa). Importantly, dimorphic cells can have connections with isomorphic cell types in only one sex (i.e. a dimorphic type with an unshared connection). Morphologically, we observed cell types that appear to be vestigial in females (Fig. 5c), and also cell types that retain the bulk of their arbors (Fig. 5d,e). However, loss of arbors does not account entirely for loss of connectivity. vMS12 and vMS11 have fewer arbors and smaller cell counts per type (vMS12, 7 female cells, 20 male cells; vMS11, 9 female, 15 male). While both retain a high degree of input, vMS12 loses virtually all relative output, but vMS11 retains some (Fig. 5c). On the other hand, although IN17A094 and IN03B024 retain their cell counts, a high degree of input, and the majority of their arbors in the female, they also have virtually no relative output (Fig. 5d). It therefore appears that a variety of cellular and circuit rearrangements can result in elimination or dampening of a cell type’s influence on the circuit. However, cells can also be retained. dMS9 receives inputs from similar cell types in males and females, but its outputs are distinct, suggesting it has retained function in females (Fig. 5e). In particular, female dMS9 targets one of the few female-specific cell types in the wing tectulum, INT2996. Further examination of INT2996 indicates that it is inhibitory, downstream of key flight steering DNs (DNa04, DNa05) and upstream of the hg (fourth axillary) steering muscles, which are centrally involved in driving song (Fig. S7) (O’Sullivan et al. 2018). These observations raise the possibility that females have acquired an inhibitory cell type that suppresses song-related activity.

### Sex Dimorphisms in the Abdominal Ganglion

Previous analyses of sexually dimorphic circuitry in the ANm were limited by damage in the previously available female VNC connectome (FANC; (Stürner et al. 2025; Azevedo et al. 2024)). Here, we have identified the full complement of sexually dimorphic and sex-specific INs in the ANm of both MANC and BANC (Fig. 6). These cell types densely arborize across the ANm, with potentially female-specific neurite extension in the anterodorsal region (Fig. 6a). The female ANm is composed of a higher proportion of dimorphic and sex-specific synapses, cells, and cell types (Fig. 6b). In both the MANC and the BANC VNC, the dimorphic and sex-specific IN types of the ANm mostly have high relative input from and output to other dimorphic and sex-specific cell types (Fig. 6c top); in contrast, isomorphic IN types mostly do not (Fig. 6c bottom). However, similar to the results presented above, a few isomorphic INs do have a fraction of input and output synapses with dimorphic and sex-specific neurons. There are documented differences present among SNs in the ANm (Rezával et al. 2012, 2014), which we have yet to characterize due to major discrepancies, in particular, in SN numbers between MANC and BANC (Bates et al. 2025). However, since sexually dimorphic and sex-specific INs of the ANm have largely remained uncharacterized, our identification presented the opportunity to understand the circuitry downstream of sexually dimorphic DNs (descending neurons) that project to the ANm.

### Abdominal Circuitry Downstream of DNp13 in Males and Females

DNp13 is one of a handful of dimorphic descending neurons with known functions; however, its complete set of connections in the ANm has not been characterized (Stürner et al. 2025).

DNp13 drives female ovipositor extrusion (which functions as both a receptivity and rejection behavior) in response to male courtship song (Wang, Wang, Forknall, Parekh, et al. 2020; Mezzera et al. 2023). In males, DNp13 has been posited to regulate song production (Stürner et al. 2025), but similar to females, male DNp13 also has projections into the ANm (Fig. 6d,f). In the male ANm, these include a set of uncharacterized visceral/circulatory efferent neurons (predicted to be neurosecretory) that target the viscera and circulation (VCs or ENs) (Bates et al. 2025; Marin et al. 2024). Outside of ANm, male DNp13 targets neurons in the wing tectulum, including vMS16, IN27X003, TN1a and vPR9; both TN1a and vPR9 have been previously noted as DNp13 targets (Stürner et al. 2025; Ding and Lillvis 2025), but our analysis shows DNp13 is not their main input (Fig. S8b). Female-only targets include both neurons in the ANm and neurons in or projecting into the wing tectulum (Fig. 6d,f). Female-specific wing targets include INXXX828_f and INXXX847_f, which may play a role in female wing flicking, another female rejection behavior that may be coupled with OE (Aranha and Vasconcelos 2018) (Fig. S8a).

With respect to DNp13’s role in ovipositor extrusion (OE) (Wang, Wang, Forknall, Parekh, et al. 2020; Mezzera et al. 2023), several MNs are targeted in the female but not male, including MNad14 and 30, which should be investigated for roles in the multiple modes of OE (Fig. 6f) <u>(Mezzera et al. 2023)</u>. In contrast, male DNp13 targets a dense network of dimorphic neurons in the ANm (Fig. 6f, magenta), which may correspond to further local regulation of ENs, as ENs are direct targets of male DNp13 (Fig. 6d and see Fig. 6h) and also arborize densely in this region. Since the effector neurons (MNs, ENs) of the ANm have yet to be matched across the male and female datasets, we grouped ANm effectors by their output nerves to better understand female versus male output to the periphery (Fig. 6g, S9, S10). The major ANm outputs to the reproductive tracts and terminalia (genitalia and analia) exit through the ANm Nerve Trunk (Taylor 1989; Cury and Axel 2023; Talsma et al. 2012). By grouping effectors, we find that female DNp13 preferentially targets ANm nerve trunk MNs, which we expect to target ovipositor extrusion muscles and perhaps other muscles (Fig. 6g). Interestingly, this female-biased DNp13 motor control runs through an isomorphic premotor layer, reflecting a trend we see elsewhere (Fig. 4b and 5a-b) for dimorphic and sex-specific neurons (they are concentrated in the pre-premotor layer rather than premotor). In contrast to the nerve trunk motor targets of female DNp13, male DNp13 preferentially targets ENs exiting via the lateral abdominal nerves. These nerves, rather than targeting the reproductive tract, combine many smaller peripheral nerves with broad, as of yet uncharacterized targets in the periphery (Power 1948; Taylor 1989; Court et al. 2020).

### Abdominal Circuitry Downstream of oviDN in Males and Females

The oviDNs (Fig. 7a) regulate egg laying in females (Wang, Wang, Forknall, Parekh, et al. 2020), though the downstream targets in the ANm have not been characterized. A recent study (Stürner et al. 2025) subdivided the population of oviDNs into 6 subtypes, only one of which (oviDNx) is proposed to be dimorphic, while the other five are female-specific. Our analysis of the oviDN neurons in the BANC supports this subdivision (Fig. 7b,c). Since males do not lay eggs, the function of oviDNx in males remains open. In addition, the function of individual oviDN subtypes in the female has not been characterized. Finally, while DNp13 and oviDN are known to control different behaviors involving the ovipositor (ovipositor extrusion (OE) versus egg laying, respectively), whether or not they share downstream partners is also not known.

We find that the dimorphic oviDNx has few shared targets in males and females (Fig. 7d). However, contrary to DNp13, oviDNx has two shared EN targets (EN00B026 and EN00B019) that are located in the ANm and exit the VNC via the nerve trunk. Previous work in *Drosophila* larvae has identified neurosecretory cells innervating the hindgut and anal sphincter in posterior VNC (Zhang et al. 2014), and in general the terminalia are innervated via the ANm nerve trunk (Taylor, 1989). Thus, these ENs may be involved in regulating sex-similar waste elimination processes.

Another EN that appears downstream of both male and female oviDNx is ab-PDF. While ab-PDF is directly downstream of oviDNx in females, it is targeted via dimorphic intermediaries in males (Fig. 7d,f, S11). ab-PDF regulates smooth muscle contraction controlling urine flow into the gut for excretion (Talsma et al. 2012). While there is slightly increased inhibition of ab-PDF via dimorphic INs in male, female oviDNx, which is excitatory, directly and strongly synapses onto ab-PDF (Fig. 7e). In females oviDNx may thus regulate urine flow via ab-PDF in conjunction with oviposition. Alternatively or additionally, pigment dispersing factor (PDF), which is released by ab-PDF neurons and stimulates renal smooth muscle contraction via volume transmission rather than local synaptic transmission, may regulate smooth muscle contraction associated with oviposition itself.

Among unshared oviDNx targets, again contrary to DNp13, there are male- and female-only connections to other ENs and very little to MNs, indicating that overall, oviDNx is likely involved in regulating peripheral processes through neurosecretory pathways rather than motor output (Marin et al. 2024). Another striking feature of oviDNx’s *downstream* circuitry is the strong recruitment of two ANs, ANXXX981 and ANXXX982 (AN_SMP_FLA_1) (Fig. 7e, S11), both of which resemble the well-characterized SAG AN (ANXXX983), which regulates post-mating behavioral changes (Feng et al. 2014) (Fig. 7f, top). Despite their morphological similarity to SAG, these two new ANs differ from SAG in their connectivity (Fig. 7f, bottom), in particular receiving no input from the sex-peptide sensory neurons (SPSNs) that drive SAG-mediated rejection behaviors (Feng et al. 2014). The intersection of these SAG-like ANs and the oviDN system in females suggests that they may be involved in oviposition-related decreases in receptivity rather than post-mating decreases in receptivity.

For the female-specific oviDNs, we find that OviDNa_a drives INXXX968_f, a female-specific octopaminergic cell type (Fig. 7g). INXXX968_f modulates a population of dimorphic and female-specific INs and ENs, which also receive overlapping input from oviDNx. This suggests that oviDNa_a is involved in female-specific modulation of circuit dynamics that are more directly driven by oviDNx. SAG appears to coordinate coactivation of these two oviDNs.

OviDNd also connects with SAG and the SAG-like ANXXX982 (Fig. 7h). It receives strong activation from SAG, and strong inhibition from ANXXX410, suggesting that these ANs are likely involved in reciprocally regulating behaviors downstream of oviDNd. OviDNd in turn targets an array of ENs, while also driving parallel feedforward inhibition of the same ENs via ANXXX982 and INXXX951_f. These ENs all exit via the abdominal trunk nerve and overlap with the EN population downstream of oviDNx and oviDNa_a, suggesting that these three oviDN subclasses all contribute differently to regulating overlapping reproductive tract functions. They are all also targeted by SAG, suggesting oviDNx, oviDNa_a and oviDNd cell types are modulated by mating state. The broad octopaminergic signalling downstream of oviDNa_a may further influence the dynamics of that state.

## Conclusion

We present a connectivity-based framework for network alignment, cell type assignment, and identification of sexually dimorphic circuits. Applied to male and female *Drosophila* ventral nerve cords, the method enables large-scale transfer of cell type annotations and the systematic identification of homologous (isomorphic), sexually dimorphic, and sex-specific neurons. By categorizing dimorphism status quantitatively from connectivity, this approach spares the need for manual morphological evaluation, simplifying the analysis pipeline and reducing potential classification bias. Using this method, we present the first comprehensive census of sex differences in intrinsic neurons (∼13,000 neurons) of the VNC. The rapid cell-typing and dimorphism discovery enabled by this approach allowed for broad assessment of sex-different circuitry of the VNC. To demonstrate the utility of our approach, we examined song-related circuits in the wing tectulum (expanding the male song circuit and characterizing the subset of these neurons present in females - an important comparison since females do not produce courtship songs) and circuits in the abdominal neuromere downstream of two dimorphic DN types known to be critical for female social and reproductive behaviors (DNp13 and oviDN). Beyond the implications for biological interpretation of connectomes, our results demonstrate that network topology alone contains sufficient information to support large-scale cell-type alignment and comparative connectomics.

As additional connectomes become available across individuals, developmental stages, and species, the primary challenge will increasingly shift from reconstruction to comparison. Future work should focus on scaling alignment algorithms to larger datasets, integrating multiple modalities of neuronal identity, and extending comparative analyses beyond sex differences.

## Methods

### Construction of ConSim from ACDC alignments

The ACDC input requires equally sized connectomes, and its output is a strict 1:1 matching of neurons in the two connectomes being compared. To satisfy the input-size constraint, disconnected padding nodes were added to the smaller graph before alignment (Fig. S1).

Because topology-only graph alignment is invariant to left-right reflection, ACDC can swap hemispheres during alignment. This ambiguity is purely orientational and can be corrected post hoc by globally exchanging left and right labels. For convenience, we fixed hemisphere orientation during alignment by adding two auxiliary nodes L and R, to each graph, with high-weight (larger than any synaptic weight) directed edges from L to R, L to left-hemisphere neurons, and R to right-hemisphere neurons. These auxiliary nodes were used only to constrain the alignment and were removed before downstream ConSim calculation and cell-type assignment.

To overcome the output limitation of strict one-to-one matching, our pipeline then uses the alignment confidence scores to construct a continuous connectivity similarity function between any pair of neurons (Fig. 1a). Specifically, for a connectome with N neurons, each neuron is embedded into a feature vector of length 2N. The first N coordinates quantify the number of input synapses received from every other neuron (presynaptic partners), and the remaining N coordinates quantify the number of output synapses sent to every other neuron (postsynaptic partners). The feature coordinates across the two connectomes are then synchronized using the 1:1 neuron-to-neuron matching produced by the ACDC algorithm. This way similarity between a query neuron **x** in one dataset and a reference neuron **y** in the other can be measured by calculating the Weighted Jaccard score (defined below) between their respective feature vectors. Importantly in this calculation, contribution of each coordinate is weighted by the continuous ACDC alignment score of the corresponding matched pair, allowing the network topology to dynamically adjust the influence of each synaptic connection (Fig. S1c). Thus, the ACDC correspondence determines which feature coordinates are compared, while the associated alignment confidence scores determine how strongly each coordinate contributes to the similarity calculation. Equipped with a similarity measure between any pair of neurons across datasets, we can identify cases where a neuron is more similar to a neuron other than its initial ACDC match and reassign it accordingly. This relaxes the rigid one-to-one correspondence imposed by graph matching and avoids artificially forcing corresponding cell types to have identical cardinalities across datasets.

### Formal Definitions

Alignment weight ω*_i_* is defined as the weighted Jaccard similarity between the σ-permuted in-and out-synaptic neighborhoods of matched node pairs *a_i_* and *b_σ(i)_*.

The similarity score between any pair of neurons *a_x_* and *b_y_* is then computed as a weighted Jaccard similarity of their directional synaptic feature vectors, scaled by coordinate-wise alignment ω_i_ of each partner.

Alignment score for σ - defined for *N* matched pairs of nodes *a_i_*, *b_σ(i)_*

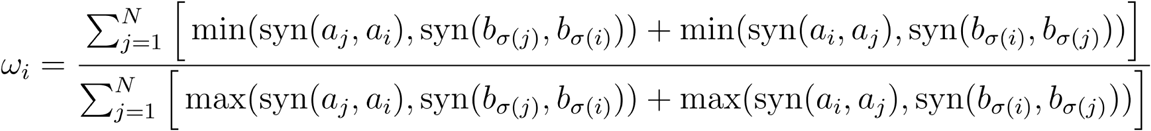

Similarity score for σ - defined for any pair of nodes *a_x_*, *b_y_*

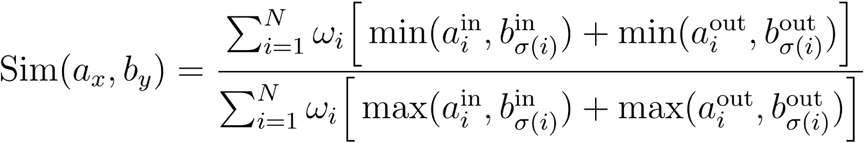

Where

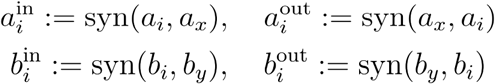

To identify dimorphisms, we compare two quantities for each neuron: its similarity to its closest cross-hemisphere twin (within the same dataset) versus its similarity to its closest cross-dataset match. For each neuron, DimScore was computed as the ratio of its highest cross-hemisphere similarity to its highest cross-dataset similarity. Cell-type DimScore was then defined as the median neuron-level DimScore across all neurons assigned to that type. This ratio provides a quantitative measure of topological variance, where a value of 1 indicates high homology. Neuron types are then classified using two discrete thresholds: a type is classified as sexually dimorphic if its median dimorphism score is between 2 and 4, and as sex-specific if the ratio exceeds 4. Thresholds of 2 and 4 were selected as simple integer cutoffs that separated previously characterized dimorphic and sex-specific neuron types while remaining conservative relative to the male–male baseline.

### Large-Scale Alignment via Data Challenge

After selecting Weighted Jaccard as the better suited similarity measure (evaluation and justification in Supplemental Methods; see also Fig. S12), we hosted a public data challenge (https://codex.flywire.ai/app/vnc_matching_challenge) attracting dozens of participants, including developers of established network alignment methods (FAQ (Vogelstein et al. 2015), SANA (Mamano and Hayes 2017)). The challenge ran from October 2024 to February 2025. Participants were provided only with the weighted directed connectivity graphs of the male and female VNCs (approximately 19,000 neurons each) and were tasked with producing a one-to-one mapping between neurons that maximized agreement of synaptic connectivity across the two connectomes. No auxiliary neuron metadata, cell-type annotations, hemisphere labels, or training correspondences were provided, making the challenge a pure graph-alignment problem based solely on network structure. A benchmark matching and scoring code were released to facilitate participation, and submissions were ranked on their ability to maximize overlap between corresponding edge weights in the aligned graphs. For simplicity, Weighted Intersection was used as the contest objective because, for fixed-size alignment problems, it induces the same ordering over alignments as Weighted Jaccard (Supplemental Methods).

### Circuit Analysis

Circuit analysis was performed with custom scripts in R and Cytoscape. Network graphs were generated with iGraph and visualized in Cytoscape via RCy3. In all network analyses, any directed cell to cell connections with <5 synapses in MANC, and <3 synapses in BANC, were omitted from the initial connectivity matrix.

Edges in single cell type comparative graphs are always input normalized; that is, they always reflect the percentage of a target cell type’s total input accounted for by the upstream cell type. We reason that, absent biophysical and functional parameters, this approach most closely approximates an upstream cell type’s influence on its downstream partner. Furthermore, MANC reconstruction recovered ∼3 times the number of synapses as in BANC VNC. Input normalization thus allows comparison of the relative input weights across the two datasets.

Edges shown in these graphs are always thresholded at the levels stated in the text except for the edges of shared inputs or outputs (middle column of nodes); as long as an edge from at least one sex reaches threshold for a shared node, the edge from the other sex is drawn regardless of weight.

## Data and Code Availability

The connectomic datasets analyzed in this study are publicly available through the FlyWire Codex (Matsliah et al. 2023) and the corresponding source publications. This includes cell type assignments and sexually dimorphic and sex-specific cell type annotations. FlyWire Codex (https://codex.flywire.ai) provides interactive web-based access to the annotations and results described in this manuscript. All analyses in this study were performed using BANC v626, MANC v1.2.1, and Male CNS v0.9. Version-specific derived data tables are archived in the accompanying GitHub repository, along with source code and data for figures (including high resolution images): https://github.com/murthylab/banc_connectivity_analysis. Source code for the ACDC alignment algorithm is available at http://bit.ly/4uxGb4q.

## Author Contributions

A.M. developed and implemented the connectome alignment methodology, designed and built the computational analysis pipeline, developed FlyWire Codex and associated tools used for data exploration, analysis, and dissemination, performed analyses, and co-wrote the manuscript. C.K.S. led the biological analyses, identified and characterized sexually dimorphic and sex-specific neuronal cell types and circuits, interpreted the results, and co-wrote the manuscript. A.S.B. and H.H.Y. contributed to cell typing and morphological verification, biological analyses, and interpretation of sexually dimorphic circuitry. A.S.B. contributed NBLAST calculations and manually reviewed connectivity matches. D.D.L. and L.K.S. invented and implemented the ACDC optimization method. B.S., J.G., S.-C.Y., K.P.W., A.T.B., R.W., and D.B. contributed annotations, proofreading, and validation. A.R.S., M.S. and C.D. managed the FlyWire community and coordinated contributor engagement. M.M. supervised the project, guided the analysis and interpretation, and co-wrote the manuscript. W.-C.A.L., H.S.S., and M.M. provided scientific guidance in development of the FlyWire-BANC dataset.

## Acknowledgments

We thank Jasper Phelps for contributions to the BANC dataset reconstruction, and Eric Perlman for data infrastructure and software support. We thank Rachel Wilson for contributions to the FlyWire-BANC connectome and support for A.S.B and H.H.Y. We thank members of the FlyWire Consortium who contributed annotations, proofreading and validation that enabled this work.

## The BANC-FlyWire Consortium Members

Alexander S. Bates, Jasper S. Phelps, Minsu Kim, Helen H. Yang, Arie Matsliah, Zaki Ajabi, Eric Perlman, Kevin M. Delgado, Mohammed Abdal Monium Osman, Christopher K. Salmon, Jay Gager, Benjamin Silverman, Sophia Renauld, Farzaan Salman, Janki Patel, Matthew F. Collie, Jingxuan Fan, Diego A. Pacheco, Yunzhi Zhao, Wenyi Zhang, Laia Serratosa Capdevila, Ruairí J. V. Roberts, Eva J. Munnelly, Nina Griggs, Helen Langley, Borja Moya-Llamas, Zuoyu Zhang, Ryan T. Maloney, Szi-chieh Yu, Amy R. Sterling, Marissa Sorek, Krzysztof Kruk, Nikitas Serafetinidis, Serene Dhawan, Finja Klemm, Paul Brooks, Ellen Lesser, Jessica M. Jones, Sara E. Pierce-Lundgren, Su-Yee Lee, Yichen Luo, Andrew P. Cook, Theresa H. McKim, Dimitrios Stasi Giakoumas, Benjamin Gorko, Justin Ellis-Joyce, Jiayi Zhang, Emily C. Kophs, Tjalda Falt, Alexa M. Negron-Morales, Austin Burke, James Hebditch, Kyle P. Willie, Ryan Willie, Sergiy Popovych, Nico Kemnitz, Dodam Ih, Kisuk Lee, Ran Lu, Akhilesh Halageri, J. Alexander Bae, Ben Jourdan, Gregory Schwartzman, Damian D. Demarest, Emily Behnke, Doug Bland, Anne Kristiansen, Jaime Skelton, Tom Stocks, Dustin Garner, Anthony Hernandez, Sandeep Kumar, Kevin C. Daly, Sven Dorkenwald, Forrest Collman, Marie P. Suver, Lisa M. Fenk, Michael J. Pankratz, Zepeng Yao, Fei Wang, Stephen J. Huston, Tomke Stürner, Gregory S. X. E. Jefferis, Katharina Eichler, Andrew M. Seeds, Stefanie Hampel, Sweta Agrawal, Tatsuo S. Okubo, Meet Zandawala, Thomas Macrina, Diane-Yayra Adjavon, Jan Funke, John C. Tuthill, Anthony Azevedo, H. Sebastian Seung, Benjamin L. de Bivort, Mala Murthy, Jan Drugowitsch, Rachel I. Wilson, Wei-Chung Allen Lee, Anna Verbe, Gabriel A. Nieves-Sanabria, Devon Jones, Zijin Huang, Sofia Pinto, Celia David, Omaris Y. De Pablo-Crespo, Emily Ye, Wolf Huetteroth, Zequan Liu & Fernando J. Figueroa Santiago

## Supplementary Methods

### Effectiveness of Similarity Measures

To evaluate effectiveness of similarity measures we considered: *Cosine* similarity, Inverse *L_1_* distance, Inverse *L_2_* distance, Weighted Intersection and Weighted Jaccard similarity. In all experiments, we generated sparse vectors for each neuron to represent its connectivity profile, including both its input and output partners (or partner types) and the corresponding synaptic weights. For connectivity vectors *u* and *v*, the evaluated similarity measures are defined as follows:

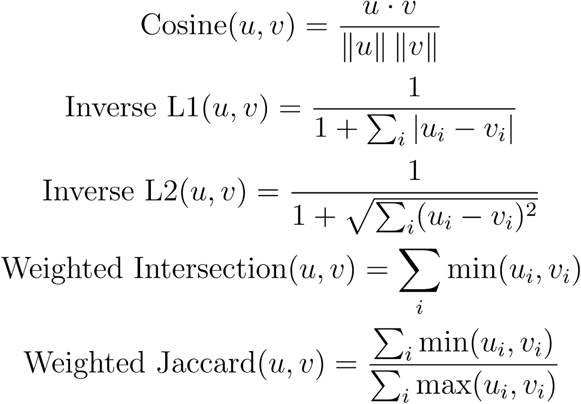

### Experiment 1: Intra-Type Retrieval Task

Our first evaluation was a retrieval task to test how well a measure can identify neurons of the same type. For each dataset, we conducted 1,000 trials, selecting a random neuron and finding its most similar partner using each of the four measures. We recorded success when the top hit matched the same neuronal type. In all 3 of the fully reconstructed datasets, Weighted Jaccard had the lowest type mismatch rate when compared to Cosine and inverse L1/L2 metrics (Fig. S12a)

### Experiment 2: Cross-Hemisphere Sub-circuit Alignment Task

Our second evaluation directly tested the ability of each measure to find the identity (*exactly correct)* alignment (graph matching) between homologous sub-circuits. We sampled 1,000 connected 10-neuron subcircuits from the left hemisphere and their ground-truth counterparts on the right. For each measure, we evaluated all *10!* possible permutations and recorded a success if the measure’s score was maximized at the identity permutation. Here too Weighted Jaccard scored highest (Fig. S12b) (for this task Weighted Jaccard and Weighted Intersection are equivalent, as well as Cosine and inverse L2 - see formal proofs in the next section).

### Equivalence of Similarity Measures for Network Alignment

For the specific task of finding the best alignment (i.e., finding the optimal permutation *P* of a neuron’s partners), two of the considered measure pairs become equivalent.

### Equivalence of Cosine Similarity and Inverse L_2_ Distance

The cosine similarity for permutation *P* is

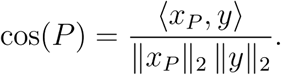

Because ‖x*_P_*‖_2_ = ‖*x*‖_2_ for all *P*, maximizing cos(*P*) is equivalent to maximizing the inner product *(^x^p, y)*

The squared Euclidean distance is

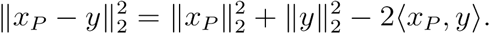

The first two terms are constant with respect to *P*, so minimizing ‖*x_P_ − y*‖_2_ (or maximizing its inverse) is also equivalent to maximizing (*x_p_,y*)

Therefore, the permutation that maximizes cosine similarity is exactly the same as the one that minimizes *L*_2_ distance (or maximizes inverse *L*_2_).

#### Equivalence of Inverse L_1_ Distance and Weighted Intersection

The *L*_1_ distance for permutation *P* is

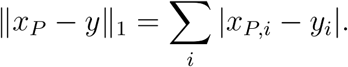

For any real numbers *u, v*,

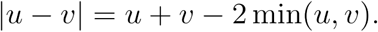

Applying this identity componentwise gives

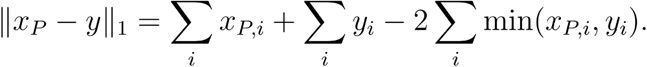

The first two sums are independent of *P*, so minimizing ‖*x_P_ − y*‖_1_ (or equivalently maximizing its inverse) is equivalent to maximizing

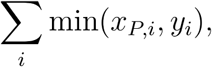

the *weighted intersection* (or coordinate-wise overlap).

Hence, the permutation that minimizes *L*_1_ distance is identical to the one that maximizes the weighted intersection score.

### Equivalence of Weighted Intersection and Weighted Jaccard

Let *A=(a_ij_*) and *B=(b_ij_*) be weighted directed adjacency matrices with *a_ij,_b_ij_*≥ 0. Let be a permutation representing a node alignment, and define the permuted matrix *b^P^_ij_ = b_P(i)P(j)_*

Define the weighted intersection

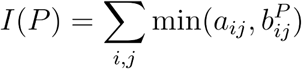

and the weighted Jaccard similarity

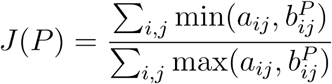

For any nonnegative *x,y* we have

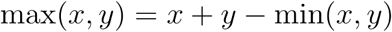

Applying this identity elementwise and summing gives

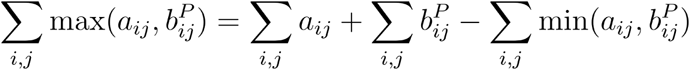

Let

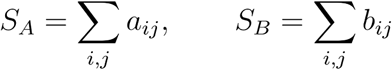

Since permutation preserves sums,

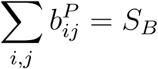

Therefore

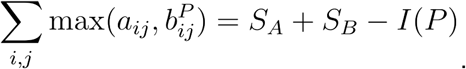

Substituting into the definition of J(*P*) yields

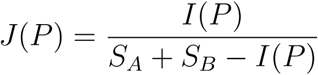

Let *C=S_A_+S_B_* which is constant with respect to *P*. Then

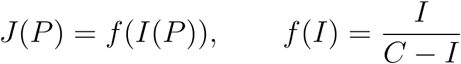

The derivative is

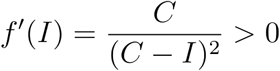

For 0≤*I*<C, so *f* is strictly increasing. Hence for any permutations *P_1_, P_2_*

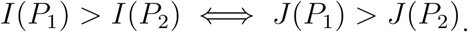

Therefore the two objectives induce the same ordering over permutations and

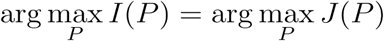

Thus maximizing weighted intersection is equivalent to maximizing weighted Jaccard similarity for adjacency matrix alignment.

## Supplementary Figures

**Fig. S1.**
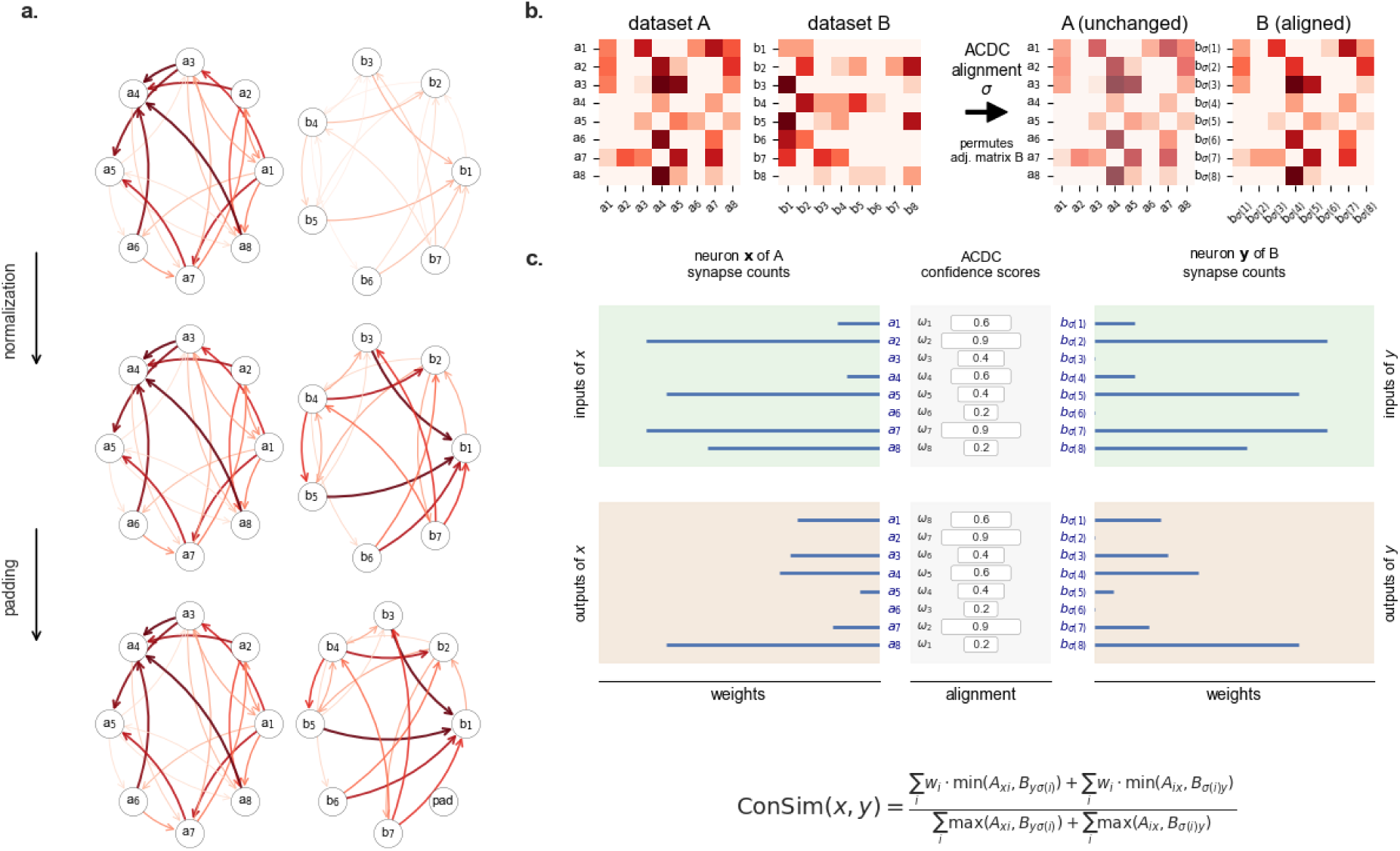
Construction of connectivity similarity scores from aligned connectomes. (a) Preprocessing of connectomes prior to alignment. Connectomes A and B are first normalized to account for differences in synaptic weights. When the connectomes contain different numbers of neurons, disconnected padding nodes are added to the smaller graph to equalize graph size. The resulting preprocessed connectomes (now compatible with ACDC) are used for subsequent alignment. **(b) Alignment using ACDC.** The adjacency matrix of connectome B is aligned to the adjacency matrix of connectome A using the ACDC graph matching algorithm. ACDC outputs a permutation matrix, σ, that reorders the neurons of B to maximize topological correspondence with A. The produced one-to-one mapping between neurons in the two connectomes also yields an alignment confidence score for each matched pair. **(c) Construction of dynamically weighted connectivity feature vectors.** Connectivity similarity is calculated between any neuron x in connectome A and any neuron y in connectome B. For each neuron, feature vectors are constructed from the numbers of input and output synapses associated with every aligned partner neuron. Each coordinate is weighted by the alignment confidence score of the corresponding matched pair, causing highly confident alignments to contribute more strongly than uncertain alignments. Connectivity similarity, ConSim(aₓ,bᵧ), is then computed as a weighted Jaccard similarity between the resulting feature vectors. The resulting score ranges from 0 to 1, with larger values indicating greater similarity in connectivity patterns.

**Fig. S2.**
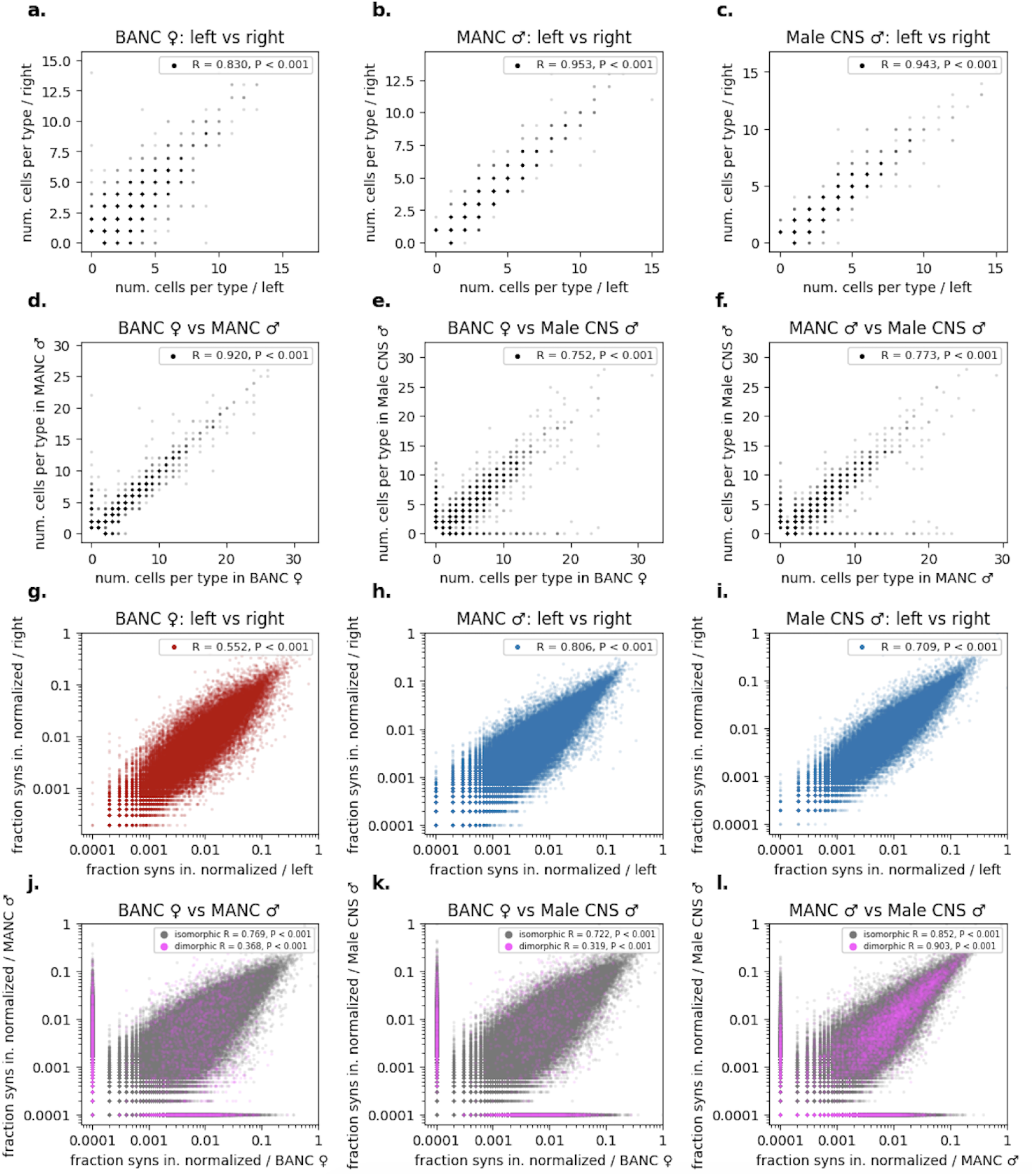
Validation of cell type assignments through bilateral symmetry and population consistency. **(a–c)** Comparing the number of cells per type in the left versus the right hemisphere for female (BANC) and male (MANC, Male CNS) datasets. High correlation indicates strong bilateral symmetry in population sizes. **(d-f)** Comparison of total cell counts per type across datasets. Significant deviations from the diagonal highlight numerically dimorphic or sex-specific cell populations. **(g–i)** Scatter plots (log scale) comparing the input-normalized fraction of synapses between cell types in the left versus the right hemisphere for BANC, MANC and Male CNS. Strong correlation along the diagonal demonstrates bilateral symmetry in synaptic weights. **(j-l)** Comparison of type-to-type synaptic fractions across sexes. Connections are stratified based on the dimorphism categories of the participating cell types.

**Fig. S3.**
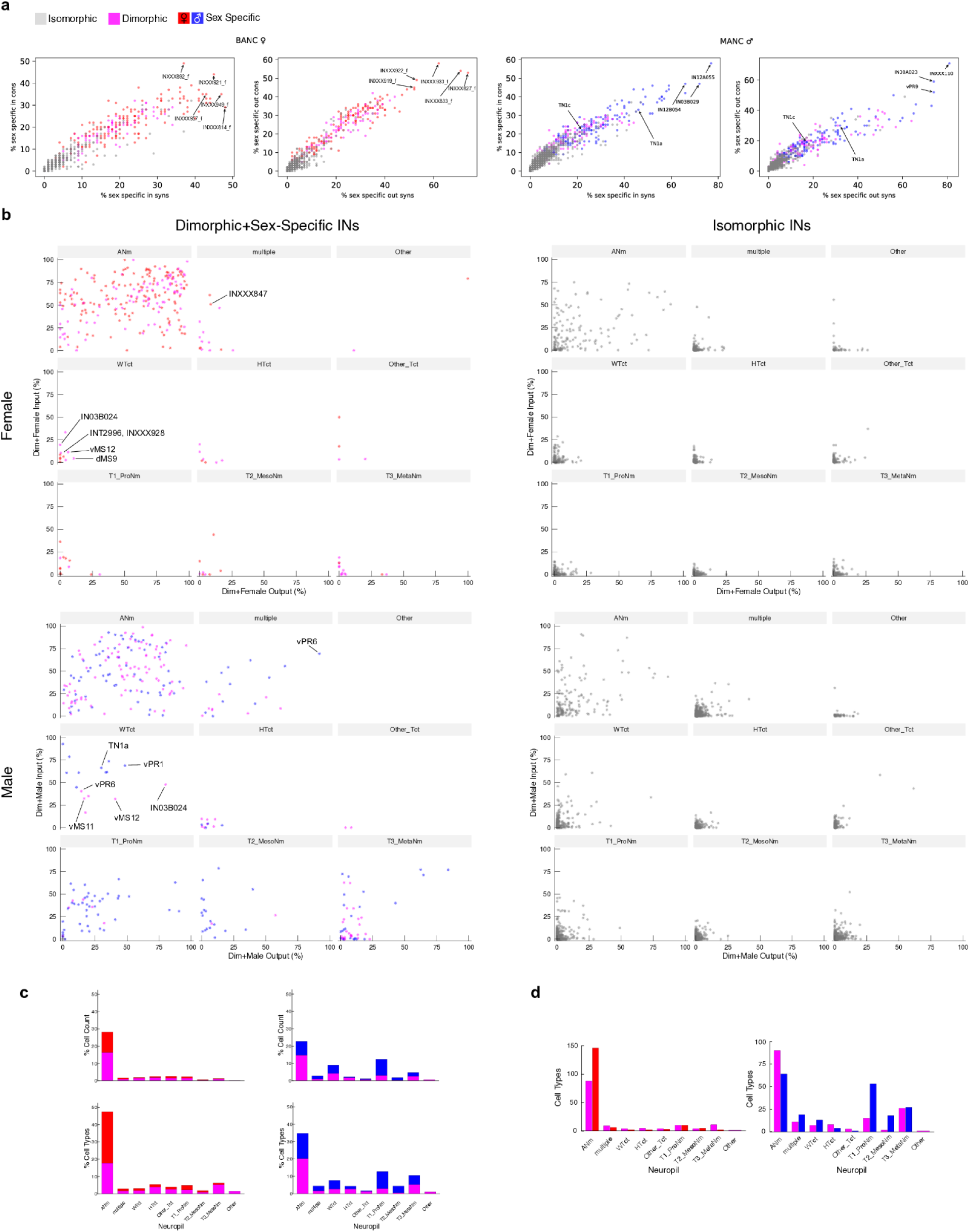
**a)** BANC VNC and MANC IN cell-type upstream connectivity (1rst and 3rd panels from left) and downstream connectivity (2nd and 4th panels) with sex-specific cells. Isomorphic cell types cluster near the origin, while dimorphic and sex-specific types exhibit significantly higher connectivity with female specific partners. **b)** Same as Fig. 3H, but considering input/output from all super classes, and faceted by neuropils. **c)** Proportions of total cells (top) and cell types (bottom) in BANC VNC (left) and MANC (left) that are dimorphic (magenta) and sex-specific (red/blue). Broken down by VNC neuropils. **d)** Absolute numbers of dimorphic and sex-specific cell types in BANC VNC and MANC.

**Fig. S4.**
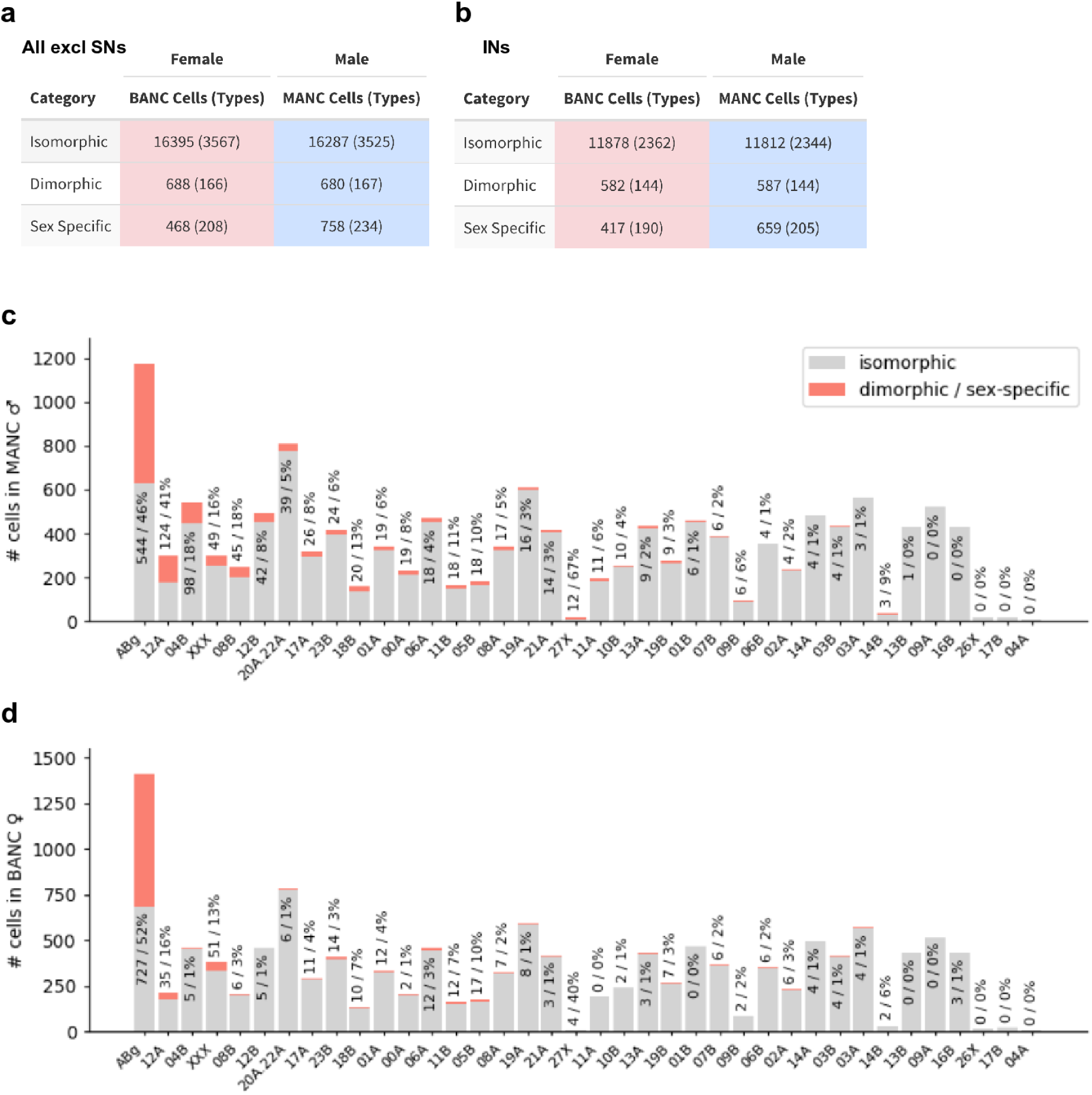
**a)** Absolute numbers of dimorphisms in all VNC neurons in BANC and MANC, excluding SNs. SNs are excluded because their numbers vary widely between BANC and MANC and matching is still incomplete. **b)** Absolute numbers of dimorphisms in all VNC INs. **(c,d)** The stacked bar charts display dimorphic and sex-specific cells (top segment) together with isomorphic cells (base segment) for each hemilineage. Neurons with >50% of their synapses in the abdominal ganglion are grouped in ABg and the vast majority of them lack hemilineage assignment in both datasets. Remaining neurons without hemilineage assignment are grouped in XXX.Text annotations denote the absolute number of dimorphic cells, followed by the percentage of the total hemilineage population that exhibits dimorphism. **(c)** Dimorphism distribution by hemilineage in the male VNC (MANC). **(d)** Dimorphism distribution by hemilineage in the female VNC (BANC).

**Fig. S5.**
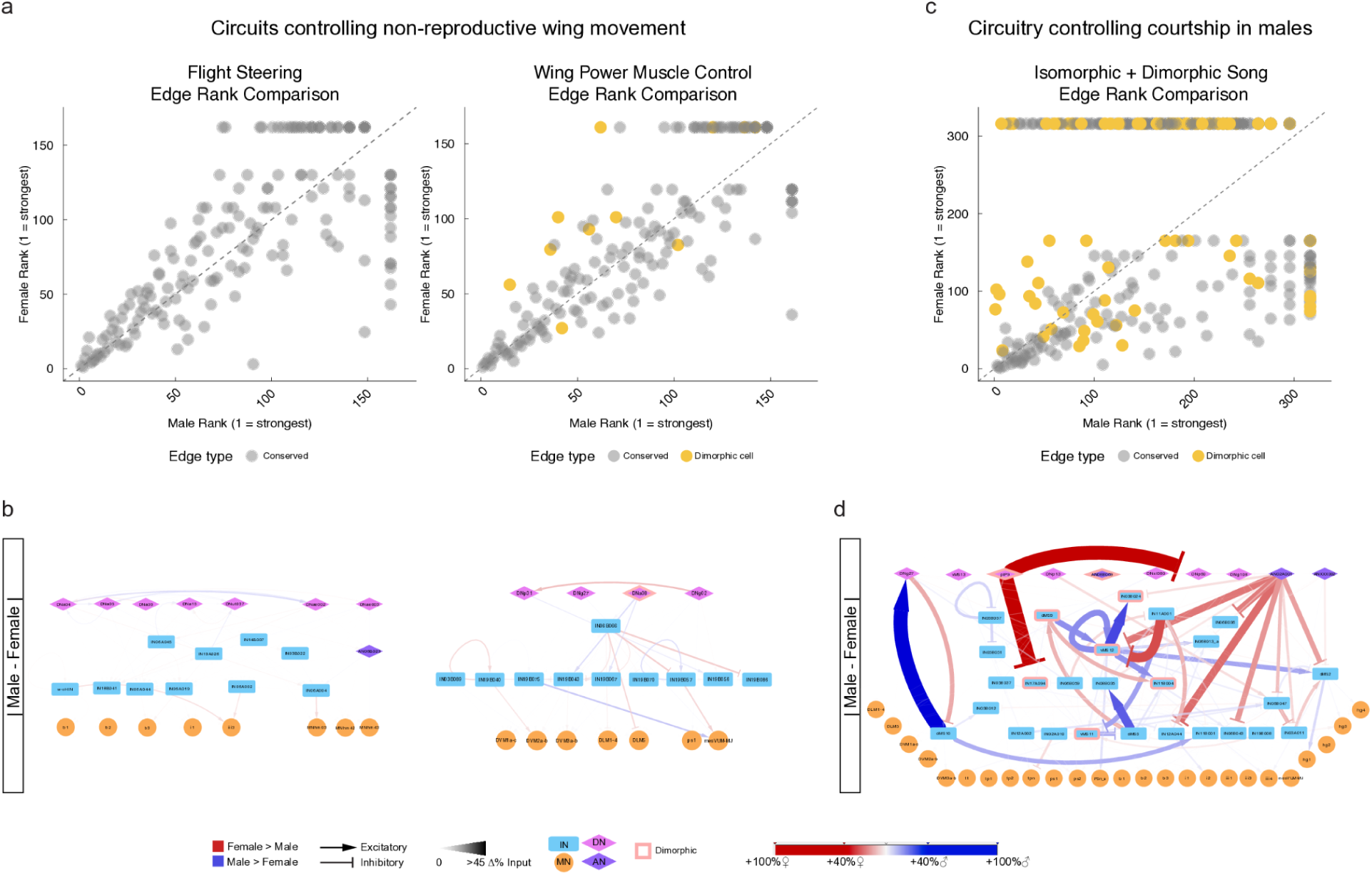
Comparative analysis of Flight and Song Circuitry. **a)** Female vs male rank correlation plots of pairwise cell type connections in the flight steering and wing power muscle motor circuits (Cheong et al., 2023) in the wing tectulum. Ranks are based on synapse counts. Points that have no connection in one sex or the other are set to the number of shared nodes + 1. Connections with shared synapse weights are set to the lowest rank. **b)** Comparative plots of the circuits in (a). Note the relative lack of cell edges that are weaker or stronger in males or females. **c)** Connection rank correlation of the sex-shared cell types of the expanded song circuit. **d)** Repetition of Fig. 5b for comparison to b.

**Fig. S6.**
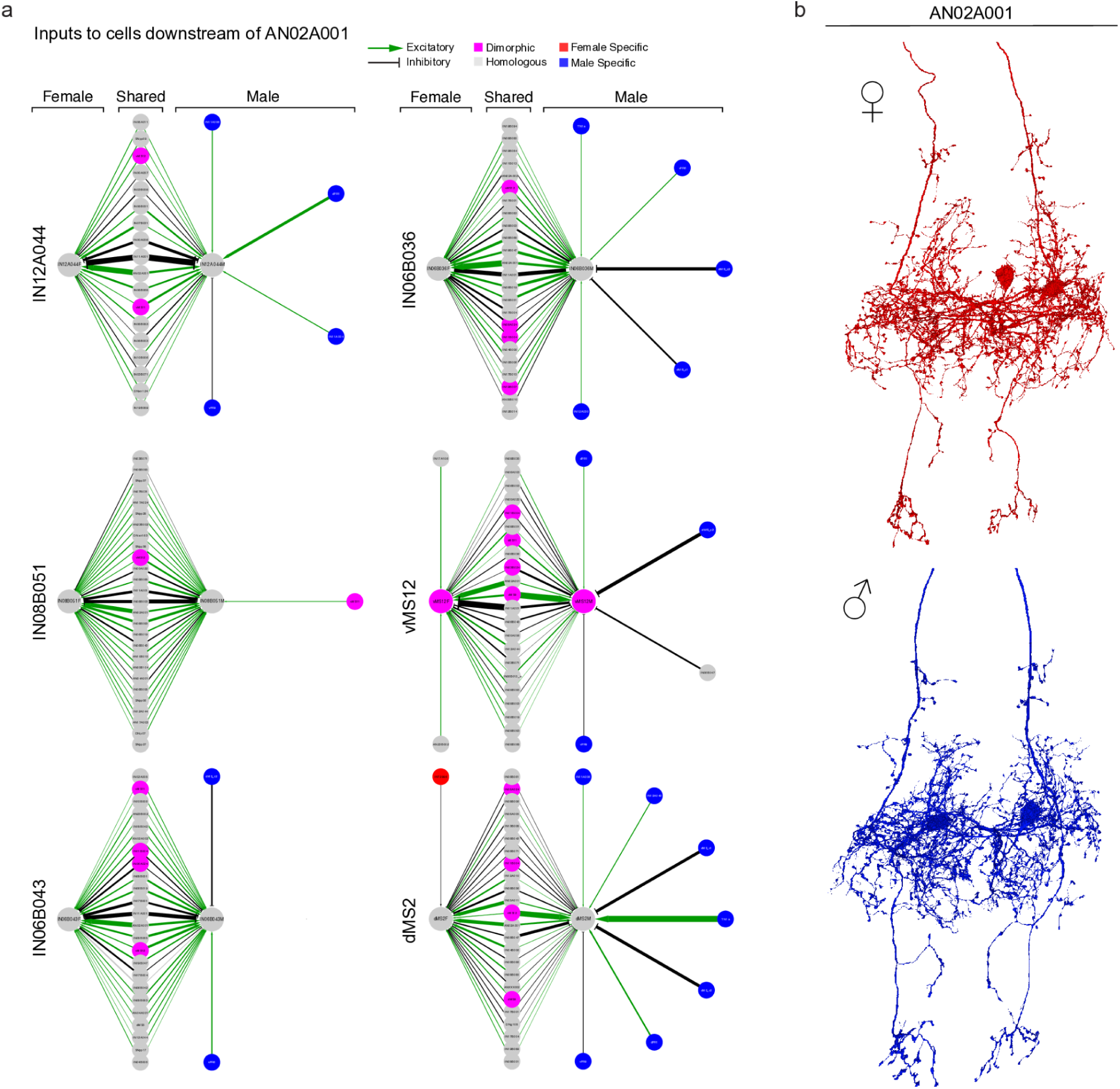
Relative increase of inhibition through AN02A001 in dimorphic female “song” neurons is due to increased male-specific inputs to AN02A001 targets. **a)** Single cell type comparative input graphs of major targets of AN02A001. Inputs were thresholded at 1% of total input to target cell. **b)** Renders of male and female AN02A001 showing sex isomorphism.

**Fig. S7.**
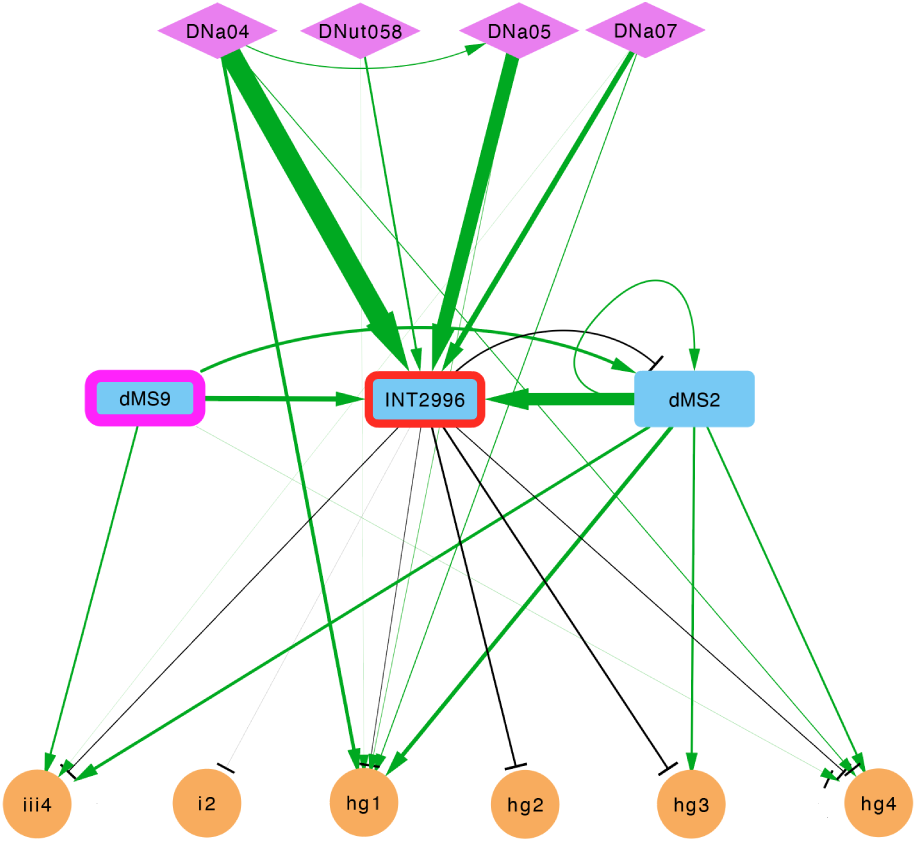
Comparative analysis of steering microcircuit; Female-specific neuron connections with song neurons. Shared motor output of INT2996 and its major inputs.

**Fig. S8.**
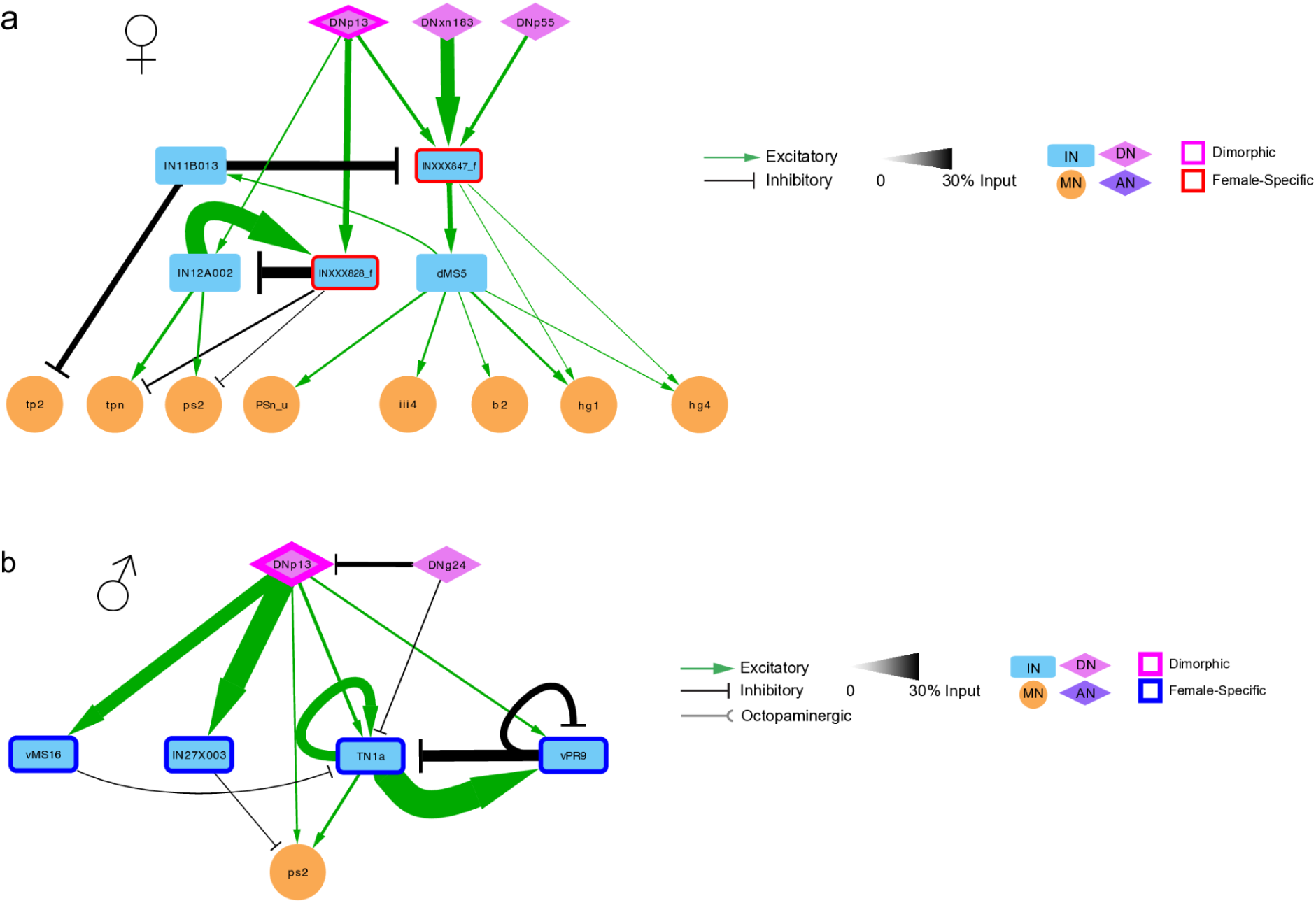
Sex-specific flight and song circuitry targeted by DNp13 in both males and females. **a)** Previous work predicted that DNp13 plays a central role in male song production (Sturner 2026, Lillvis 2024). This was based on the percentage of DNp13 output synapses targeting cell types known to be involved in song production (e.g., in MANC, DNp13’s top target is TN1a, receiving 5.8% of DNp13 output synapses; 1.7% of DNp13 output synapses are to vPR9). However, DNp13 has many targets, and when considering the proportion of its targets’ input synapses that DNp13 accounts for, it only ranks as TN1a’s twelfth strongest input (2.5% input) and vPR9’s thirteenth (1.5% of input), suggesting it does not exert may not exert major influence over these cells. By contrast, pIP10, the major descending regulator of courtship song, provides the strongest descending input to TN1a and vPR9 (28.4% and 9.4% of input respectively) On the other hand, DNp13 accounts for a large proportion of the input to vMS16 (14%) and IN27X003 (24.6%), both male-specific GABAergic cells targeting wing and song neurons. Future functional studies should examine DNp13’s influence on song through these cell types as well. Wing motor neuron ps2 is included as it is the only direct MN target of DNp13 in the wing. **b)** DNp13 targets two female-specific neurons in the wing tectulum. INXXX847_f is predicted to excite PSn_u, iii4, b2, hg1 and hg4 via dMS5 (all of which are implicated in male courtship song (Lillvis et al. 2024; Shirangi et al. 2016; Clemens et al. 2018; O’Sullivan et al. 2018)) and INXXX828_f is predicted to inhibit tpn and ps2 (not known to be associated with song) both directly and via IN12A002. Combining their female-specificity and DNp13’s role in ovipositor extrusion rejection behavior (Wang, Wang, Forknall, Parekh, et al. 2020) suggests that these female-specific neurons may be involved in driving mating-associated wing behaviors such as wing flick rejection (Spieth 1974; Aranha and Vasconcelos 2018).

**Fig. S9.**
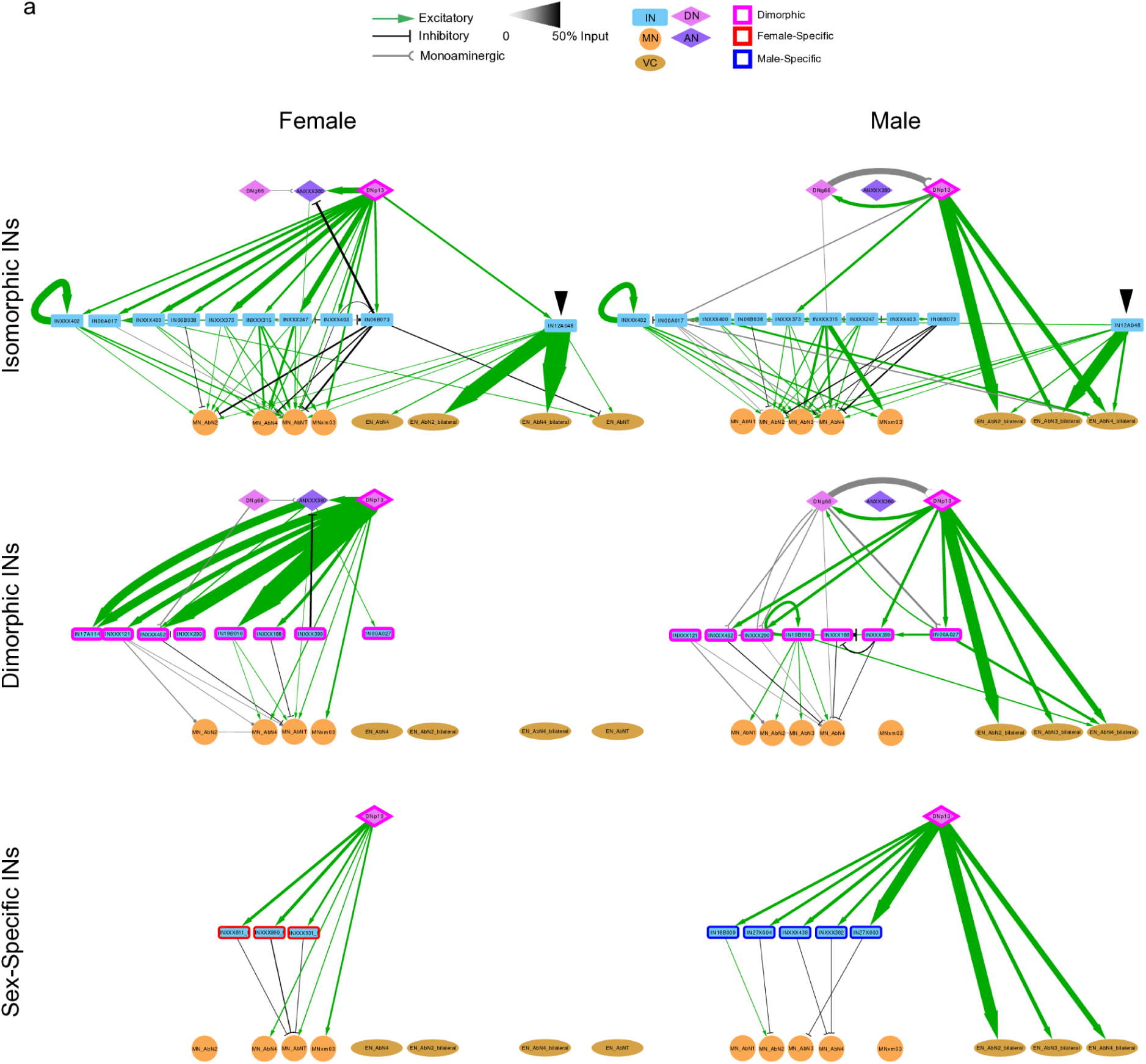
Isomorphic, dimorphic and sex-specfic pre-effector targets of DNp13. Downstream IN and effector (MN and EN) neuron targets of female (left) and male (right) DNp13. Isomorphic and dimorphic cell types above a 1% input threshold in females OR males are included in both sexes; sex-specific cells (bottom row) included in that sex only. Arrowhead indicates loss of direct DNp13 input to IN12A048

**Fig. S10.**
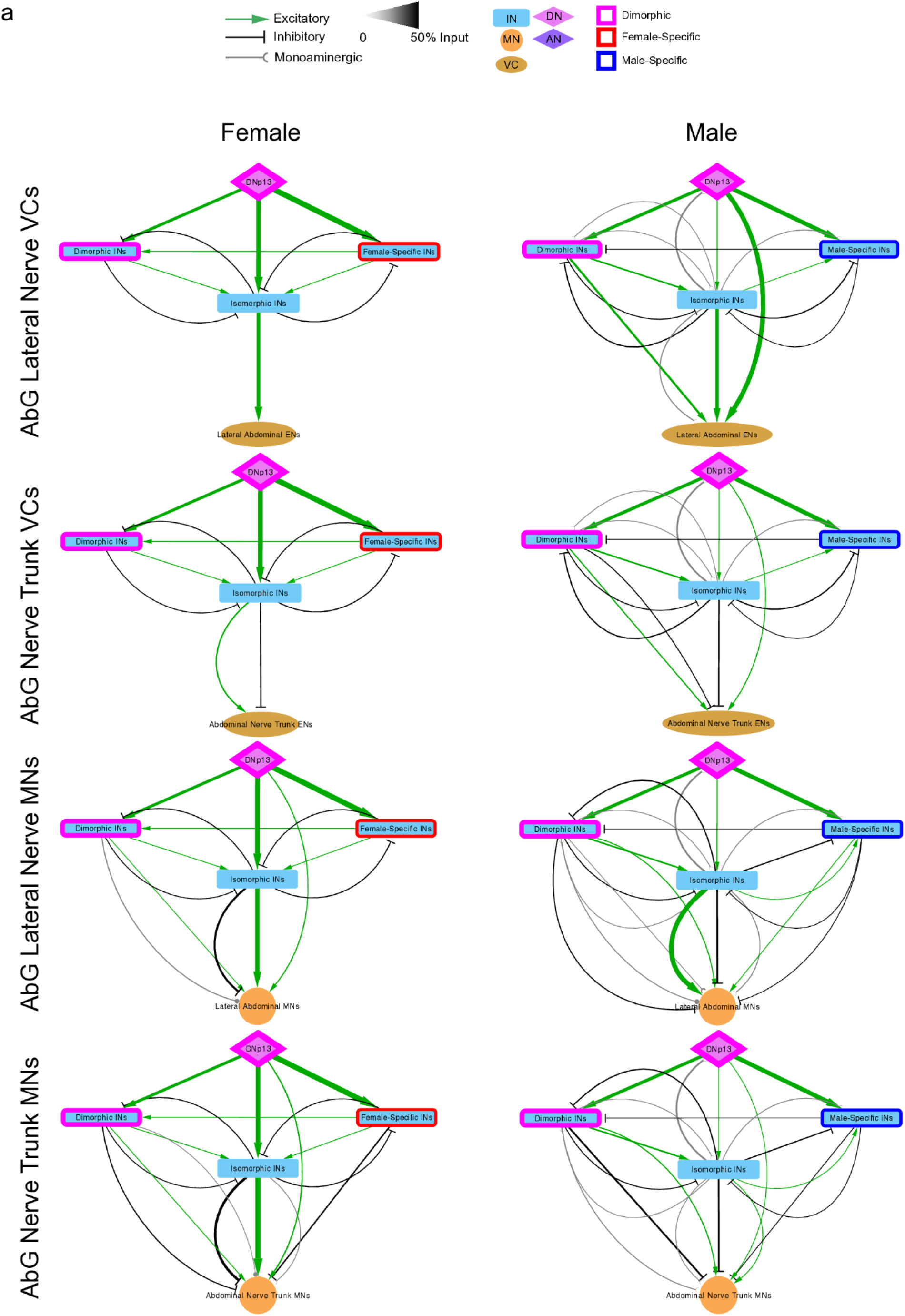
Coarse comparative analysis of male and female DNp13 effector targets. Proportional input DNp13 and its directly downstream isomorphic, dimorphic and sex-specific partners onto ENs exiting the lateral or trunk ANm nerves (top two rows) and MNs exiting the lateral or trunk ANm nerves (bottom two rows.)

**Fig. S11.**
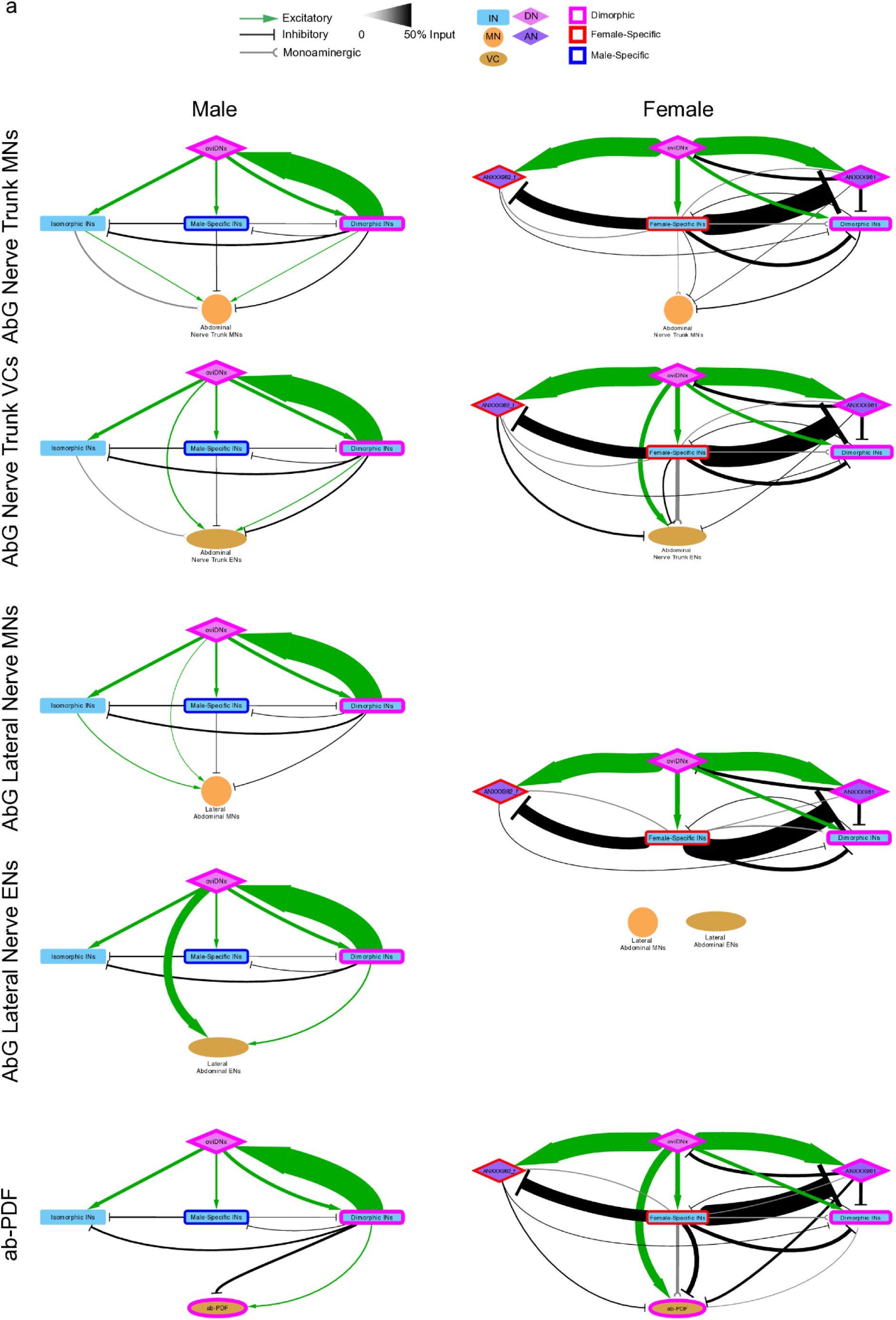
Coarse comparative analysis of male and female oviDNx effector targets. Proportional input male (left) and female (right) oviDNx and its directly downstream isomorphic, dimorphic and sex-specific partners make onto ENs exiting the lateral or trunk ANm nerves (top two rows); MNs exiting the lateral or trunk ANm nerves (third and fourth rows); and ab-PDF (bottom row).

**Fig. S12.**
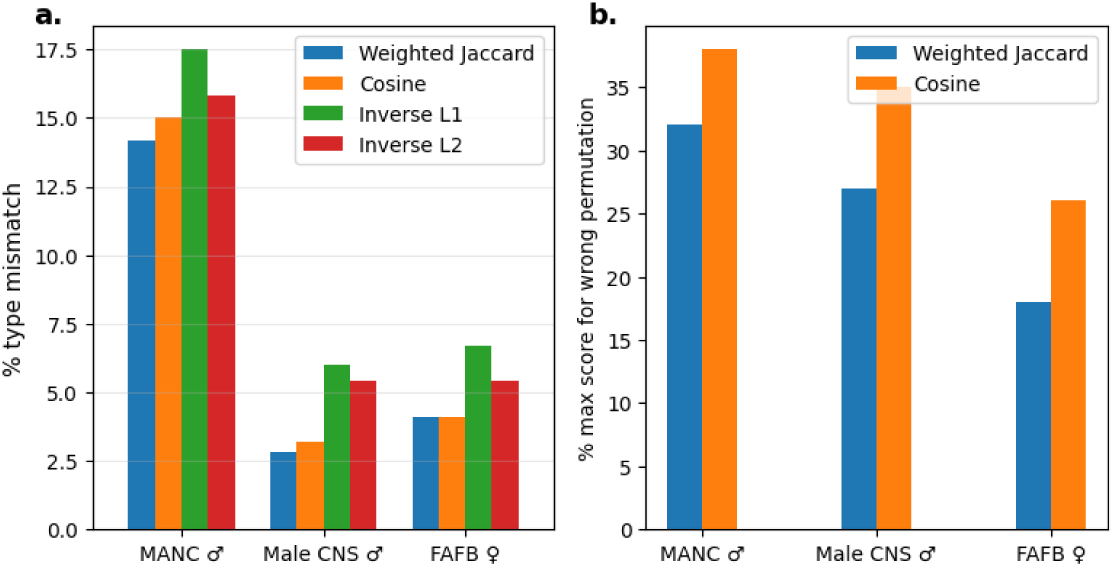
Evaluating similarity measures. We assessed the accuracy and effectiveness of different similarity metrics for neuron connectivity. **(a)** Intra-dataset neuron retrieval: For each dataset and measure, we performed 1,000 trials. In each trial, a random query neuron (excluding untyped and singleton cells) was matched to its closest neighbor using feature vectors composed of input and output synapse counts grouped by partner types. Error rates are reported as the percentage of trials where the closest match was not the same cell type as the query. **(b)** Evaluating similarity measures for cross-hemisphere sub-circuit alignment: To test the ability of different metrics to identify the exact topological correspondence between homologous networks, we sampled 1,000 pairs of isomorphic, connected 10-neuron sub-circuits across the left and right hemispheres. To ensure an unambiguous ground truth, these sub-circuits were composed exclusively of singleton cell types (neurons with exactly one instance per hemisphere). For each sub-circuit pair, we evaluated all possible mappings (10! permutations). We compared Weighted Jaccard and Cosine similarity; as demonstrated in the Methods, for the specific mathematical task of optimizing permutations, Weighted Jaccard is equivalent to inverse L1 distance and Weighted Intersection, while Cosine similarity is equivalent to inverse L2 distance. Error rates represent the percentage of the 1,000 trials where the similarity metric failed to be globally maximized by the known ground-truth alignment. Lower error percentages indicate superior alignment accuracy.

